# An Improved Genome Assembly of *Azadirachta indica* A. Juss

**DOI:** 10.1101/033290

**Authors:** Neeraja M. Krishnan, Prachi Jain, Saurabh Gupta, Arun K. Hariharan, Binay Panda

**Affiliations:** Ganit Labs, Bio-IT Centre, Institute of Bioinformatics and Applied Biotechnology, Biotech Park, Electronics City Phase I, Bangalore 560100, India; Strand Life Sciences, Bellary Road, Hebbal, Bangalore 560024, India

**Keywords:** PASA, paired-end, mate-pair, *FDFT1*, *SQLE*, Platanus, PacBio, error-correction, LoRDEC, neem, assembly, training-set, gene prediction, gene structure, genome, transcriptome

## Abstract

Neem (*Azadirachta indica* A. Juss.), an evergreen tree of the *Meliaceae* family, is known for its medicinal, cosmetic, pesticidal and insecticidal properties. We had previously sequenced and published the draft genome of the plant, using mainly short read sequencing data. In this report, we present an improved genome assembly generated using additional short reads from Illumina and long reads from Pacific Biosciences SMRT sequencer. We assembled short reads and error-corrected long reads using Platanus, an assembler designed to perform well for heterozygous genomes. The updated genome assembly (v2.0) yielded 3- and 3.5-fold increase in N50 and N75, respectively; 2.6-fold decrease in the total number of scaffolds; 1.25-fold increase in the number of valid transcriptome alignments; 13.4-fold less mis-assembly and 1.85-fold increase in the percentage repeat, over the earlier assembly (v1.0). The current assembly also maps better to the genes known to be involved in the terpenoid biosynthesis pathway. Together, the data represents an improved assembly of the *A. indica* genome.

The raw data described in this manuscript are submitted to the NCBI Short Read Archive under the accession numbers SRX1074131, SRX1074132, SRX1074133, and SRX1074134 (SRP013453).

## Introduction

High-throughput sequencing platforms, especially those based on short-read technology, have enabled sequencing of many plant genomes (Michael and Jackson, 2013). This has substantially improved our understanding of genome organization, evolution and complexity in different plant species. However, most first generation genome assemblies are draft and incomplete assemblies. The correctness and accuracy of genome assembly depends on the length of the sequencing reads, errors generated during sequencing and the accuracy of the computational tools (assemblers and downstream annotation pipelines) used. Additionally, most genome assemblers are not suitable to assemble genomes of heterozygous plants, a characteristic feature of most plants in the wild (Kajitani, Toshimoto, et al., 2014). Draft assemblies often bear significant gaps and, errors, yielding less accurate gene predictions and annotations. This is compounded by the usage of incomplete training-sets with gene prediction algorithms and absence of a representative transcriptome that can correctly anchor to the genome. Therefore, it is imperative to improve the quality of draft genome assemblies with the help of longer reads using genome assemblers tailored to handle heterozygosity, and make gene predictions using updated training-sets and gene annotations using combinatorial approaches not fully reliant on sequence similarity such as BLAST.

Neem (*Azadirachta indica* A. Juss.), belonging to the order *Rutales,* family *Meliaceae,* is an important woody angiosperm, given its many medicinal and agrochemical uses. We had previously sequenced and reported the draft genome and five organ-specific transcriptomes (Krishnan, Pattnaik, et al., 2011, Krishnan, Pattnaik, et al., 2012) of the neem tree. The neem genome was the 38^th^ plant genome to be sequenced (Michael and Jackson, 2013). The genome assembly was generated using short paired-end reads (76 bases or shorter) from Illumina GAIIx with a first generation genome assembler, SOAPdenovo (Li, Zhu, et al., 2010). This was followed by genome annotation and gene prediction analysis, analysis of repeat elements, phylogenetic analysis and gene expression studies (Krishnan, Pattnaik, et al., 2012). In the current report, we have improved the quality of the neem genome assembly by using [a] additional long-insert libraries from Illumina Hiseq, [b] long reads from a third generation sequencer by Pacific Biosciences (PacBio), [c] LoRDEC (Salmela and Rivals, 2014), an algorithm that takes short reads from Illumina and uses those to correct errors in the PacBio reads, and [d] assembling the genome with short and error-corrected long reads using Platanus (Kajitani, Toshimoto, et al., 2014) which is better suited to assemble heterozygous genomes. We re-assembled all five organ-specific RNA libraries into a pooled representative transcriptome, using Trinity (Grabherr, Haas, et al., 2011, Haas, Papanicolaou, et al., 2013), and employed the Program to Assemble Spliced Alignments (PASA, Haas, Delcher, et al., 2003) to benchmark the completeness of previous (v1.0), intermediate, and current (v2.0) genome assemblies based on their mappability to this transcriptome. We also performed gene prediction analyses with GlimmerHMM (Majoros, Pertea, et al., 2004, v3.0.4) using updated training-sets from Citrus species, which were found to be evolutionarily closer to neem by our earlier phylogenetic analyses (Krishnan, Pattnaik, et al., 2012). Building on our draft assembly, here, we present data on different assembly parameters, accuracy, gaps, gene predictions and the total repeat content as evidence towards an improved neem genome assembly.

## Materials and Methods

### Assembly

In addition to the Illumina read libraries used for assembling the previously published draft neem genome (Krishnan, Pattnaik et al. 2012), four more libraries were used for updating the assembly. We included reads from three Illumina mate-pair (with insert sizes 4kb, 6kb and 10kb) and one PacBio (average read length >2kb, varying up to 17.64kb) libraries. Details of all libraries used are presented in Supplementary Table 1.

We pre-processed all the libraries as follows. In the case of Illumina libraries, exact read duplicates were removed using the ‘in *silico* normalization’ utility from Trinity. For PacBio, reads were error-corrected using LoRDEC v0.4.1 based on the two paired-end Illumina libraries (Supplementary Table 1). K-mers ranging from 19 to 36 were tested for error-correction.

We made an effort to assemble intact PacBio reads following error-correction using the PacBioToCA (Koren, Schatz, et al., 2012) pipeline and Celera WGS assembler v7.1 (Myers, Sutton, et al., 2000). However, this process was CPU- and RAM-intensive, and also resulted in a sub-optimal assembly (data not shown). We, therefore, converted the PacBio reads, with and without error-correction, into Illumina-like paired-end reads (read lengths of 100 bases and average insert size of 350 bases) using SInC’s read generator (Pattnaik, Gupta, et al., 2014), which could be easily assembled using SOAPdenovo, SOAPdenovo2 (Luo, Liu, et al., 2012) and Platanus. Converting PacBio reads to Illumina-like reads did not nullify the advantage of the long reads, in terms of contiguity (Supplementary File 1).

We produced 13 intermediate assemblies (Supplementary Table 2) for quality comparison, as follows:

a. re-assembly of the published version using SOAPdenovo with Illumina short reads (R.S1/v1.0)
b. assembly using additional Illumina libraries using SOAPdenovo2 (S2.DUP)
c. assembly of all Illumina duplicate-removed libraries using SOAPdenovo2 (S2)
d. assembly, using SOAPdenovo2, of all Illumina duplicate-removed libraries along with the error-corrected PacBio reads (S2.ecPB.21 and S2.ecPB.32, using kmers 21 and 32, respectively)
e. assembly using Platanus of all Illumina duplicate-removed libraries alone (P), or along with either the error-corrected PacBio reads using 19- (P.ecPB.19), 21- (P.ecPB.21), 32- (P.ecPB.32) and 36-mers (P.ecPB.36), or along with uncorrected PacBio reads (P.ucPB)
f. assembly and gap closing, using Platanus, of all Illumina duplicate-removed libraries and the PacBio library with (P.ecPB.32.gc/v2.0; kmer = 32) or without (P.ucPB.gc) error-correction.

All assembly QCs were performed using QUAST v2.3 (Gurevich, Saveliev, et al., 2013). The assembly NG50 was estimated assuming the neem genome size as 364Mb (Krishnan, Pattnaik, et al., 2012). We refer to the R.S1 assembly as v1.0 (previous) and the P.ecPB.32.gc assembly as the improved v2.0 (current), in our comparisons statistics below.

### Assembly mapping to transcriptome using PASA

PASA r20140417 was used to compare and evaluate all the assemblies. The representative neem transcriptome was assembled *de novo* using Trinity v2.0.6 with five tissue-specific published RNA-seq libraries. This transcriptome was mapped to various genome assemblies using PASA and the numbers and lengths of valid alignments, failed alignments, and transcript assemblies were compared. In addition, the numbers and lengths of exon-only regions of the valid alignments were also extracted and compared across the assemblies.

### Gene prediction using GlimmerHMM

GlimmerHMM was used for benchmarking the assemblies. We created training-sets based on *Citrus sinensis* and *Citrus clementina* (genes.gff3 files downloaded from http://phytozome.jgi.doe.gov/pz/portal.html), and used the inbuilt *Arabidopsis thaliana* training-set to predict genes and gene structures in the neem assemblies. Both citrus species were used here since they were found to be the evolutionarily closest to neem, among sequenced species (Krishnan, Pattnaik, et al., 2012).

### Repeat analyses

RepeatModeler v1.0.8 (Smit and Hubley, 1989), employing Repeat Scout, Tandem Repeat Finder and Recon modules, was used to construct a library of novel repeats entirely based on the neem genome. Other tools such as LTR_finder v1.0.5 (Xu and Wang, 2007), TransposonPSI v08222010 (Haas, 2007–10) and MITE-hunter v11-2011 (Han and Wessler, 2010), were used to identify Long Terminal Repeats (LTRs), retro-transposons, and Miniature Inverted repeat Transposable Elements (MITEs), respectively. The neem genome assembly was masked using RepeatMasker v4.0.5 (Smit and Hubley, 1989) with all these repeats and the updated plant (*Viridiplantae*) libraries from Repbase (Kapitonov and Jurka, 2008), to estimate the non-redundant genomic repeat content. This was further classified using the RepeatClassifier module of RepeatModeler.

*Identification of* FDFT1 *and* SQLE *gene structures across assemblies*

We obtained the transcript sequences corresponding to *FDFT1* and *SQLE* genes in *C. clementina* from KEGG (Kanehisa and Goto, 2000; Kanehisa, Goto, et al. 2014), and created a database of these sequences using the makeblastdb utility in the BLAST package v2.2.29 (Altschul, Gish, et al. 1990). These genes belong to the sesqui- and tri-terpenoid biosynthesis pathways, involved in the synthesis of the commercially important compound, azadirachtin, and hence were chosen for comparative analyses here. The neem transcriptome was mapped against the database using BLAST with an Expect (E) value threshold of 0.001. The mapped neem transcripts were traced to their PASA alignments in various genome assemblies. In cases where the identified transcripts for the same reference gene aligned to multiple neem scaffolds, consensus exon-intron structures were inferred individually for each scaffold, and the one agreeing best with the *C. clementina* gene structure was considered. The gene structures for all assemblies were plotted along with the corresponding gene structure in *C. clementina* using ‘Structure Draw’ (http://www.bioinformatics.uni-muenster.de/tools/strdraw/index.hbi). Regions of gaps (N’s) in the assembly were highlighted in red.

All scripts used in the assembly, QC, evaluation, genome-to-transcriptome mapping, gene prediction and repeat analyses pipeline are presented under Supplementary File 2.

## Results

### Quality comparison across all versions of neem genome assembly

We compared the correctness and completeness of all the assembly versions based on three measures:

1. Assembly statistics using QUAST
2. Metrics from transcriptome-to-assembly alignment using PASA
3. Gene and gene-structure prediction based on three different training-sets using GlimmerHMM

The first measure strictly quantifies the completeness of the assembly, while the middle one mainly quantifies the correctness of the assembly, and its completeness to the extent that the draft transcriptome is complete, and the last measure quantifies the completeness of the assembly, but also its correctness, with the assumption that the genes and gene structures in the organisms used as a training-sets are present, as is, in the neem genome. Detailed metrics from all the benchmarking tools are provided in Supplementary Table 2.

### Comparison of assembly statistics

Overall, assembly statistics improved with Platanus over SOAPdenovo or SOAPdenovo2 (Figure 1 and Supplementary Table 2), with the best assembly (v2.0) produced by Platanus using a combination of all duplicate-removed Illumina read libraries and error-corrected (*kmer* = 32) PacBio library in all the three stages - assembly, scaffolding and gap-closing. The scaffold numbers and the assembly size here were reduced by 2.6- and 3-fold, respectively, over those from the earlier draft assembly (v1.0; Figure 1). The assembly using uncorrected PacBio reads, in combination with Illumina libraries (P.ucPB), resulted in the longest scaffold (12,211,325 bases). However, other important quality metrics were compromised for this assembly. N50 and N75 were highest for Platanus assembly using all Illumina-only reads (P; 4,002,232 and 1,489,583 bases, respectively). The v2.0 assembly revealed a 13.4-fold reduction in gaps over the v1.0 assembly (an average of 5414.21 Ns per 100kb, Figure 1A) and a 2.26-fold lowered NG50. Incidentally, the NG50 for the Platanus assembly using Illumina-only reads (P; 1,587,838 bases) was comparable to that using SOAPdenovo (v1.0; 1,663,167 bases). Almost 60% of each assembly was covered at 5X when PacBio reads were assembled, along with Illumina read libraries, using SOAPdenovo2 or Platanus (Supplementary Table 2).

**Figure 1.**
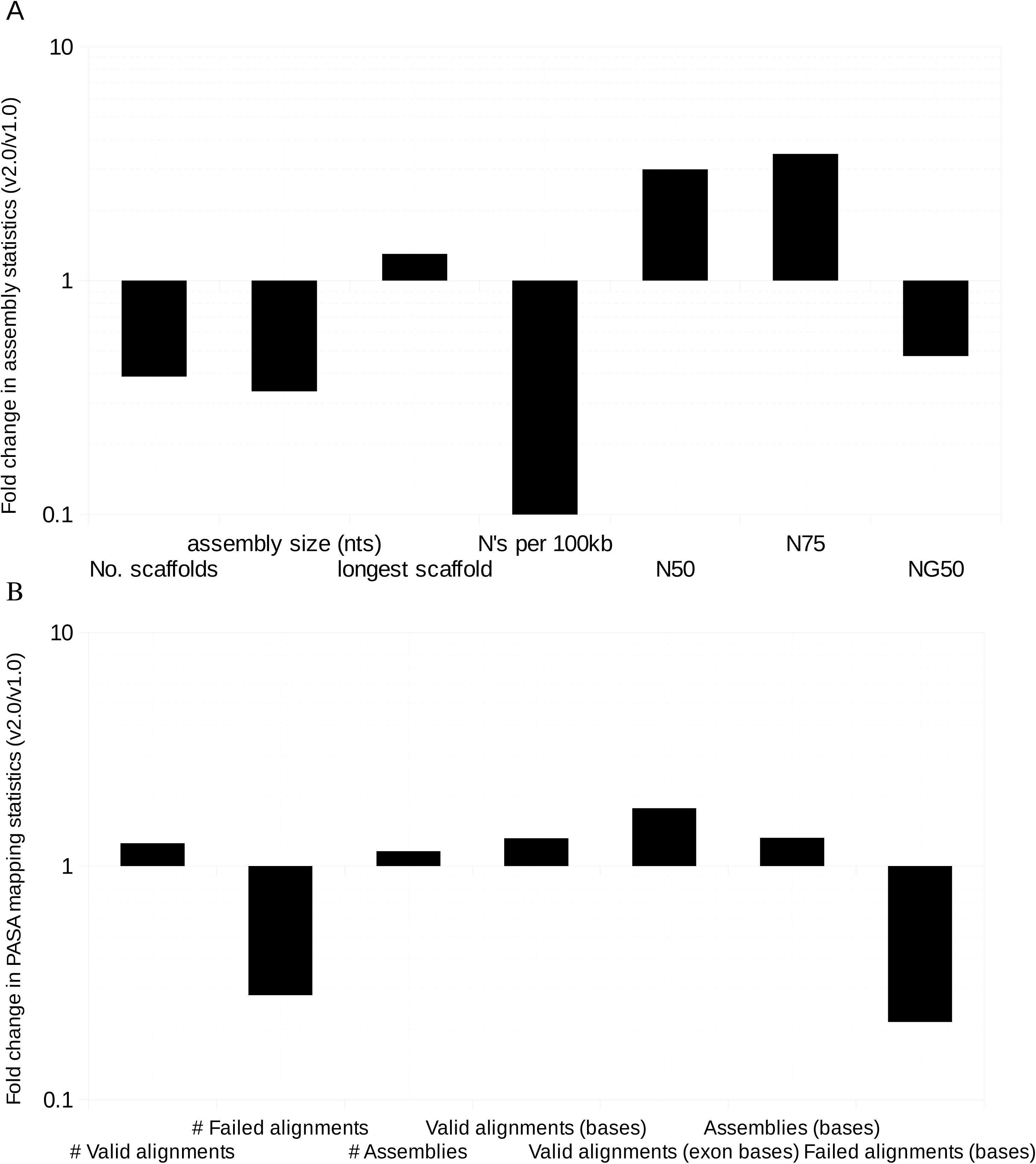
Improvements (fold change between current, v2.0, over the previous, v1.0, assembly) in various A: assembly statistics and B: PASA mapping statistics. The Y-axis is plotted on a logarithmic scale and the minor grids conform to uniform intervals on positive and negative Y axis.

### Comparison of transcriptome-to-genome alignment metrics

The numbers and cumulative lengths of all valid alignments and PASA assemblies were highest at 77,635 and 61,292, ~100Mb and ~99Mb, respectively, for the v2.0 assembly (Supplementary Table 2). The cumulative size of valid exonic alignments was also highest at ~48Mb for this assembly, and the corresponding numbers and lengths of all failed alignments were least at 6,584 and ~32Mb, respectively (Supplementary Table 2). The overall valid alignments increased 1.25-fold, and the ones in exons increased by 1.95-fold for the updated (v2.0) assembly over the old one (v1.0) (Figure 1B). Failed alignments went down by 3.5- and 5.9-fold in number and cumulative size, respectively (Figure 1B).

### Comparison of predicted genes

We found the highest number of predicted genes and exons, using training-sets from any of the three organisms (*A. thaliana, C. sinensis, C. clementina*), with the v2.0 assembly (Supplementary Table 2 and Figure 2). The cumulative length of all predicted genes was highest for this assembly (68,723,917 bases) when *A. thaliana* was used as the training-set. When *Citrus* species were used as training-sets, however, the v1.0 assembly resulted in the highest cumulative predicted gene lengths (473,787,912 and 431,305,649 bases, respectively, with *C. sinensis and C. clementina*). The predicted gene lengths were comparable between both the assemblies after excluding gaps, suggesting this to be mostly a result of mis-assembly (Figure 2).

**Figure 2.**
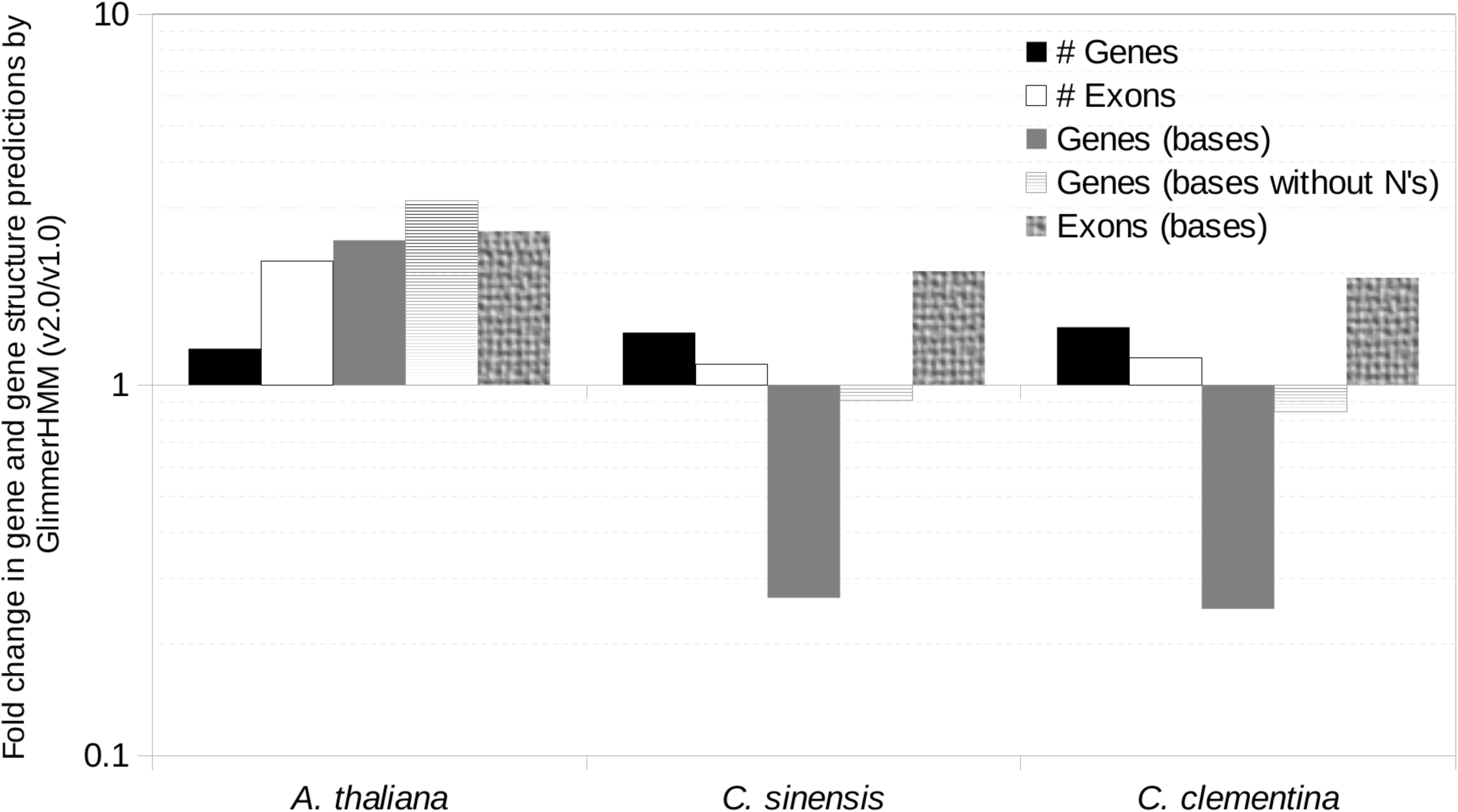
Improvements (fold change between current, v2.0, over the previous, v1.0, assembly) in the numbers (#s) and sizes (bases) of gene and exon predictions from GlimmerHMM. The Y-axis is plotted on a logarithmic scale and the minor grids conform to uniform intervals on positive and negative Y axis.

We found an abundance of smaller (< 100 bases) mRNAs and exons in gene predictions in the v1.0 assembly, especially with *Citrus* training-sets, which were substantially reduced in the v2.0 assembly (Figure 3). In contrast, the longer mRNAs were more abundant in the latter assembly, with Citrus training-sets, an indication of improvement in the assembly.

**Figure 3.**
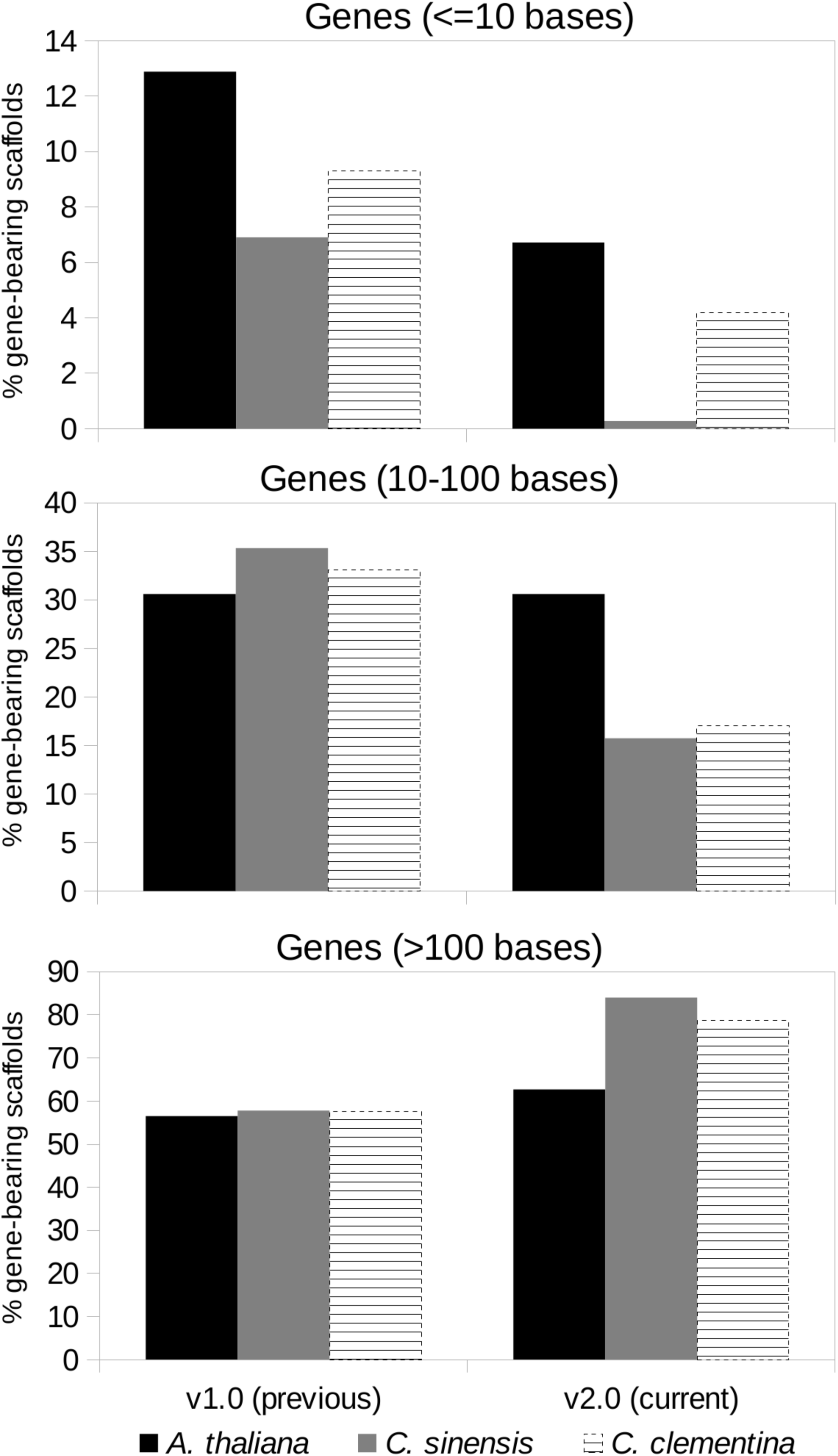
Proportion (%) of gene-bearing scaffolds/contigs with gene predictions of lengths <10 bases, 10-100 bases, and >100 bases, for *A. thaliana, C. sinensis* and *C. clementina* training sets.

### *Comparison of gene structures of* FDFT1 *and* SQLE *across various assemblies*

In order to demonstrate the biological significance of the improved assembly, we used *FDFT1* and *SQLE* genes, two important genes involved in the sesqui- and tri-terpenoid biosynthesis pathways. We observed that the gene structures of *FDFT1* and *SQLE* were more complete and accurate in the improved v2.0 assembly when compared to the v1.0 assembly (Figure 4 and Supplementary Figure 2). Using Platanus alone, and augmenting the libraries with additional short Illumina mate-pair libraries yielded a better *FDFT1* gene structure. Similarly, using Platanus as an assembler along with pre-and post-processing yielded a better assembly of the multi-isoform *SQLE* gene.

**Figure 4.**
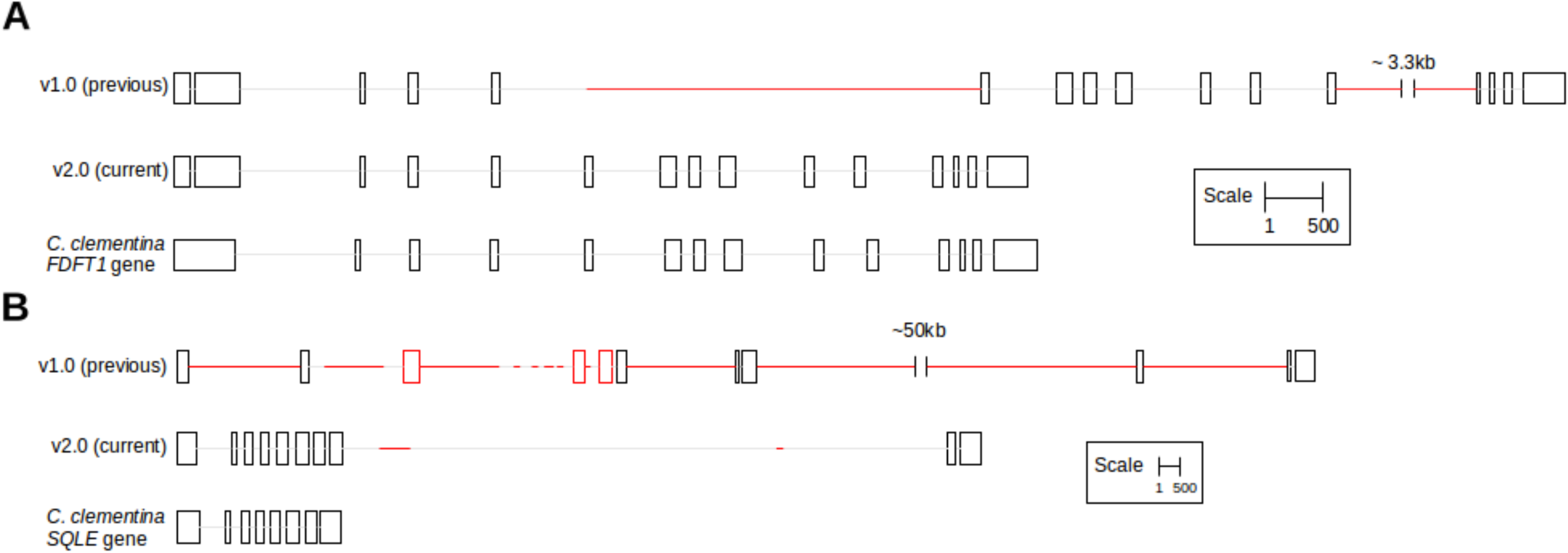
Comparison of v1.0 and v2.0 assemblies for A: *FDFT1* and B: *SQLE* genes. The *FDFT1* and *SQLE* transcripts from *C. clementina* were mapped to the representative Trinity-assembled *A. indica* transcriptome using NCBI BLAST (*E*-value 0.001). The transcripts were traced to their neem genomic scaffold mappings from PASA, in order to extract the exon-intron structures of the corresponding genes. In the figures, boxes and lines denote exons and introns, respectively, and the red regions denote gaps in the assemblies. The scales are different for *FDFT1* and *SQLE* and are, therefore, indicated individually.

### Estimation of repeat content

The non-redundant repeat content was estimated to be 54,375,206 bases (24.15% of v2.0), which is higher than the 47,427,034 bases reported earlier (13.03% of v1.0). We further classified the repeats into distinct classes, as shown in Supplementary Table 3.

## Discussion

Here, we report an improved genome assembly of *A. indica* and provide quantitative evidence on various parameters in support of the improved assembly. The current assembly benefits from using additional Illumina mate-pair reads and long reads from PacBio, a third generation sequencing platform. Additionally, we have used Platanus, a tool designed to assemble heterozygous genomes, such as that of neem (Supplementary Figure 1), better, and an algorithm that uses short reads to correct the errors in long reads. Finally, we have used updated and near complete training-sets from closely related plant species to predict gene structures, and an equally enriched and updated repeat library to predict repeat sequences in the neem genome.

In our study, we employed PASA and GlimmerHMM to benchmark the assemblies, both of which have their limitations in the current context. PASA assumes that the transcriptome is free of mis-assembly errors. The caveat with GlimmerHMM, is that the gaps and errors in the genomic assembly extends to the predictions (Figure 2). We found that the number of gene predictions decreased across assemblies, post-redundancy removal using cd-HIT-EST (Li and Godzik, 2006). Additionally, the gene predictions are only as good as the training-sets used. Presence of a large number of very short, possibly spurious, exons in the *C. clementina* training-set manifested in a large number of similar predictions in the neem assembly (Figure 3). However, as expected, either these did not align to the neem transcriptome, or a large fraction of those that aligned did not meet the validity criteria set by PASA, suggesting incorrect predictions. This implied a larger number of gene predictions not to be an indicator of correct or complete assembly in neem. Instead, integration of results from multiple tools, preferably using additional information from orthogonal high-throughput platforms such as RNA-seq, and experimental validation, offered better benchmarking.

The presence of duplicate reads may give false assurance to the assembler in terms of artificially inflated read depth. Hence, removing exact read duplicates reduces mis-assemblies. Interestingly, we found that the assembly with SOAPdenovo2, after duplicate removal (S2), displayed worse statistics, but much improved transcriptome-genome mappings using PASA (Supplementary Table 2). SOAPdenovo, using fewer Illumina libraries, and without a duplicate removal step (v1.0), also displayed sub-optimal assembly statistics but a good NG50 number (Figure 1). This, most likely, is due to an abundance of gaps in the assembly, inflating the assembly size. Incidentally, the NG50 numbers for assemblies using libraries from the same platform were comparable (Supplementary Table 2). Such observations caution against deriving conclusions regarding best assembly based solely on assembly statistics tools, such as QUAST.

Exploring the finer details of individual genomic features, instead of macro-level statistics like NG50, may provide a better estimate of the improvement in the assembly quality, as exemplified by the improved assembly of *FDFT1* and *SQLE* genes in the improved neem assembly. Relying solely on sequence similarity-based approaches for gene identification can result in incomplete and/or inaccurate structural annotations. Using BLAST against *C. clementina* transcripts, with a stringent *E*-value threshold of 0.001, identified only portions of the *FDFT1* and *SQLE* genes in our scaffolds, making us falsely deduce that we had assembled only certain exons from these genes. This would particularly be true for structurally conserved genes, which have few very important, and, therefore, conserved domains. In such genes, variable domains might not have significant sequence homology to the reference database(s) that include sequences from other species, causing the genes to not be annotated in their entirety. Therefore, our approach, of using the sequence similarity between *C. clementina* and neem transcripts to trace back the entire gene sequence, structure and combining both reference- and *de* novo-based identification techniques, is a better one (Figure 4).

In conclusion, genome assemblies need to be updated continuously by implementing accurate computational algorithms and supplementing with experimental evidence to obtain error-free and near complete assemblies. The process of obtaining accurate genome assembly is a dynamic and continuous process that needs to be undertaken, in our opinion, by groups or communities that have produced the first draft sequence of various genomes. This will facilitate research in genomics and create public resources to understand gene structure and function in plants better.

## Acknowledgements

Research presented in this article is funded by Department of Electronics and Information Technology, Government of India (Ref No:18(4)/2010-E-Infra., 31-03-2010) and Department of IT, BT and ST, Government of Karnataka, India (Ref No:3451-00-090-2-22).

## Author Contributions

BP: Overall planning, conception and design of the study, data interpretation and manuscript writing; NMK: Conception, analysis and interpretation of data, manuscript writing; PJ and SG: Analysis and interpretation of data, manuscript writing; and AKH: Sequencing data production.

## Supplementary Figure Legends

**Supplementary Figure 1.**
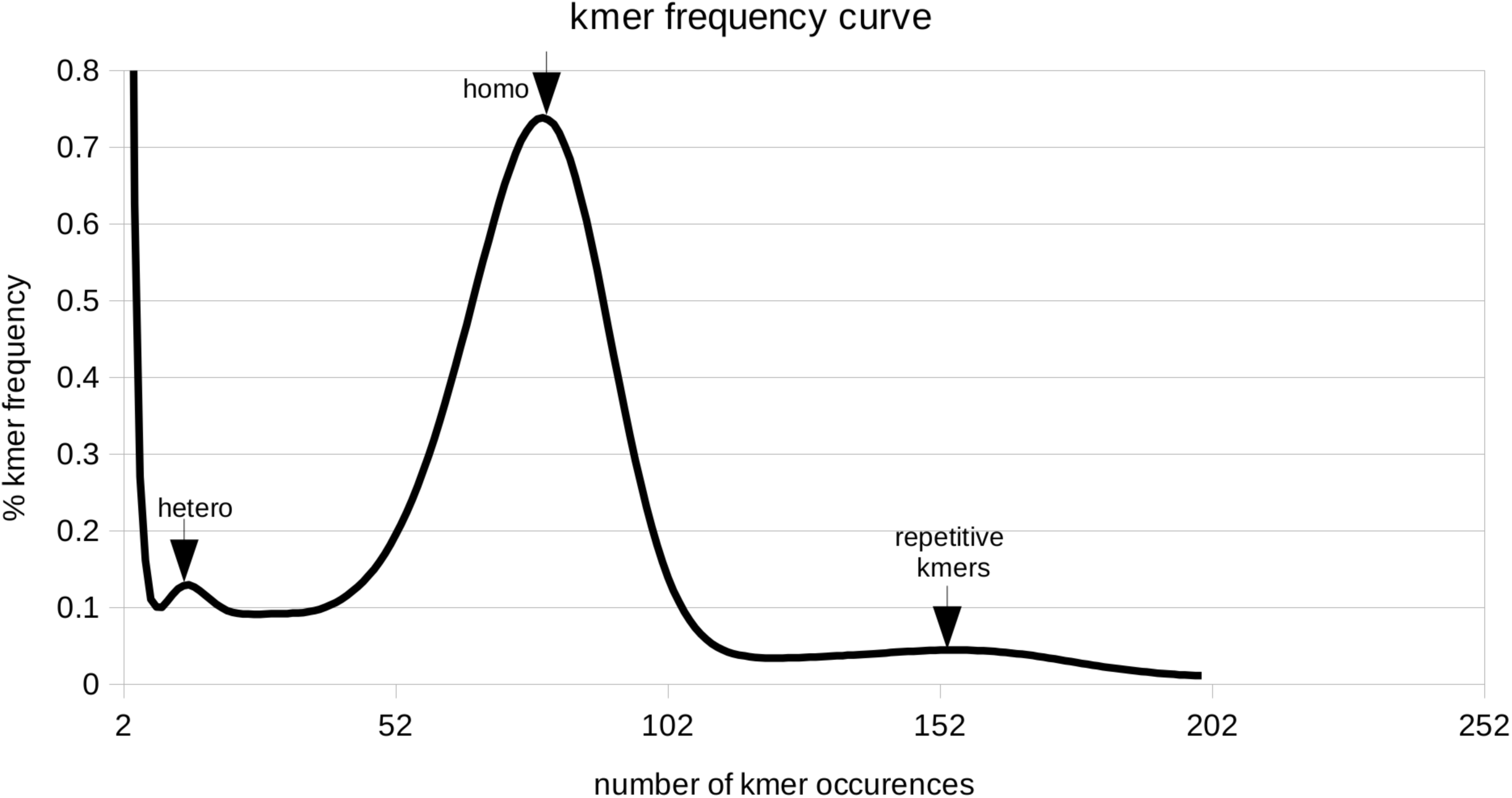
kmer frequency curve. The frequency (%) of 17-mers is plotted as a function of the number of times they occur across paired-end libraries. The peaks for heterozygous, homozygous and repetitive kmers are highlighted by arrows.

**Supplementary Figure 2.**
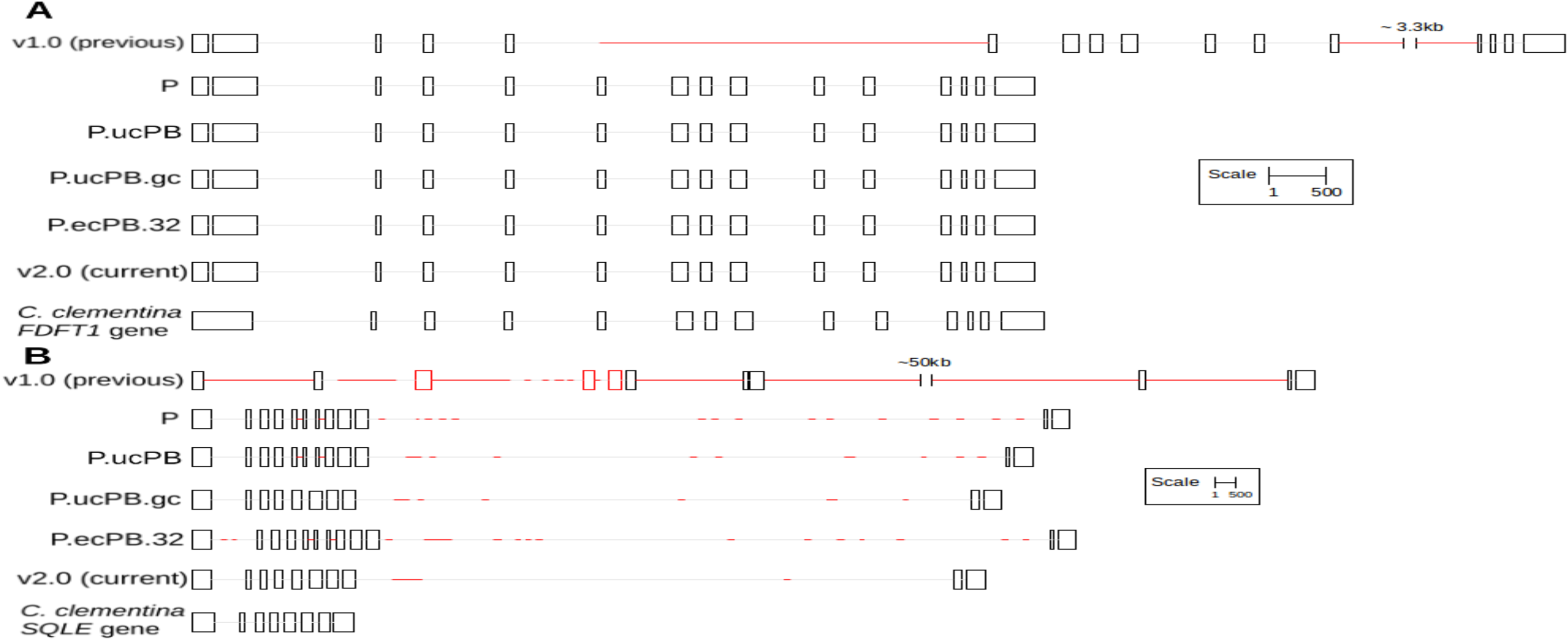
Comparison across assemblies for A: *FDFT1* and B: *SQLE* genes. The *FDFT1* and *SQLE* transcripts from *C. clementina* were mapped to the representative Trinity-assembled *A. indica* transcriptome using NCBI BLAST (*E*-value 0.001). The transcripts were traced to their neem genomic scaffold mappings from PASA, in order to extract the exon-intron structures of the corresponding genes. In the figures, boxes and lines denote exons and introns, respectively, and the red regions denote gaps in the assemblies. The scales are different for *FDFT1* and *SQLE* and are, therefore, indicated individually.

**Supplementary Table 1.** Deatils of sequencing libraries. PE: short-insert paired end, MP: long-insert mate pair libraries.

| Lib no. | Read length | Insert size | Type | Used for earlier published assembly (v1.0)* using SOAPdenovo | Used for v2.0 and intermediate assemblies |
| --- | --- | --- | --- | --- | --- |
| 1 | 76 | 350 | Illumina PE | ✓ | ✓ |
| 2 | 76 | 350 | Illumina PE | ✓ | ✓ |
| 3 | 36 | 1500 | Illumina MP | ✓ | ✓ |
| 4 | 36 | 3000 | Illumina MP | ✓ | ✓ |
| 5 | 36 | 10000 | Illumina MP | ✓ | ✓ |
| 6 | 100 | 4000 | Illumina MP |  | ✓ |
| 7 | 100 | 6000 | Illumina MP |  | ✓ |
| 8 | 100 | 10000 | Illumina MP |  | ✓ |
| 9 | Variable (mean >2Kbp, longest 17.64Kbp) | NA | PacBio |  | ✓ |

\* The original assembly contained both Ion Torrent and Sanger libraries with very low coverage (0.5X and 0.001X, respectively) which were excluded from our current assembly.

**Supplementary File 1.**
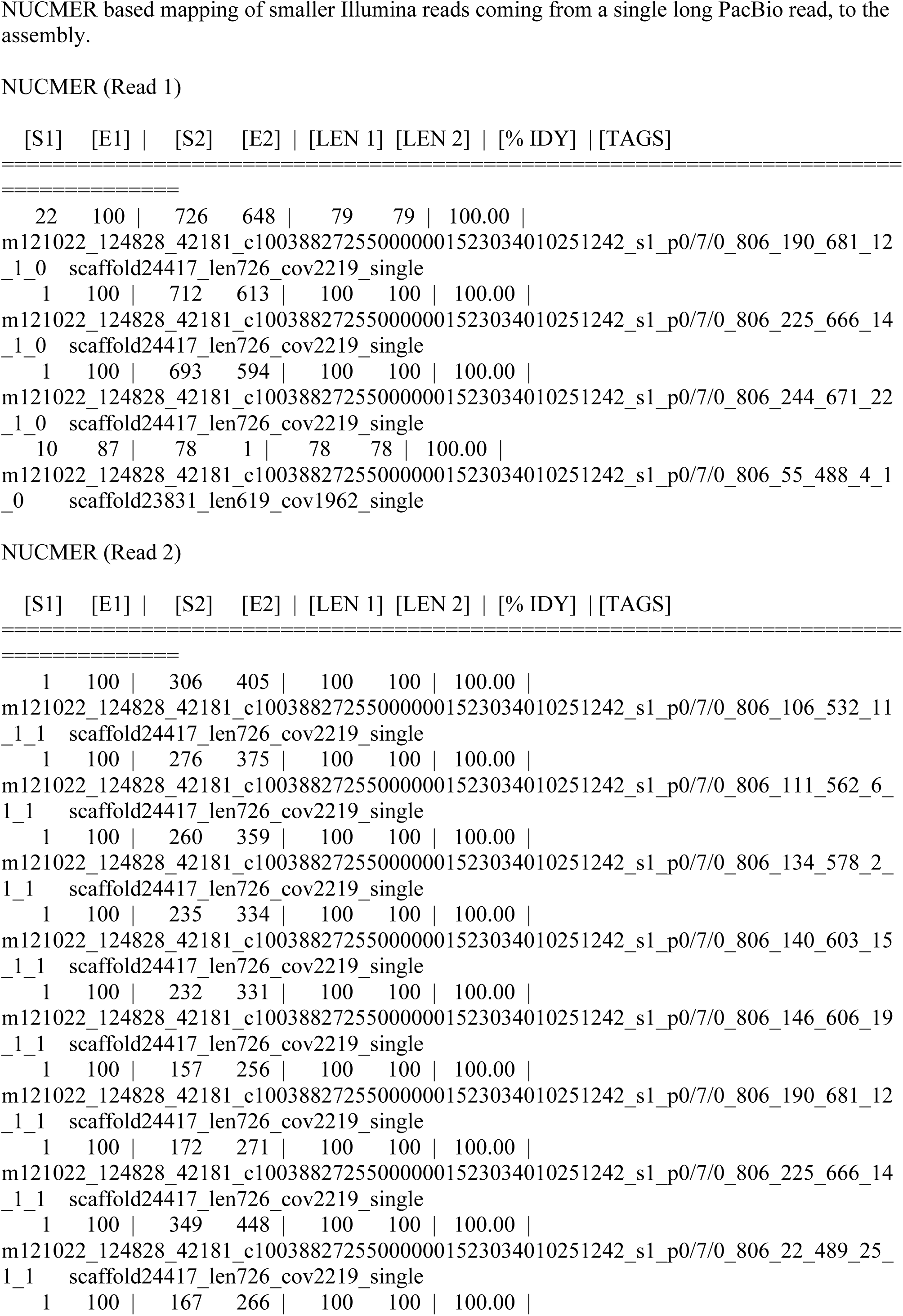
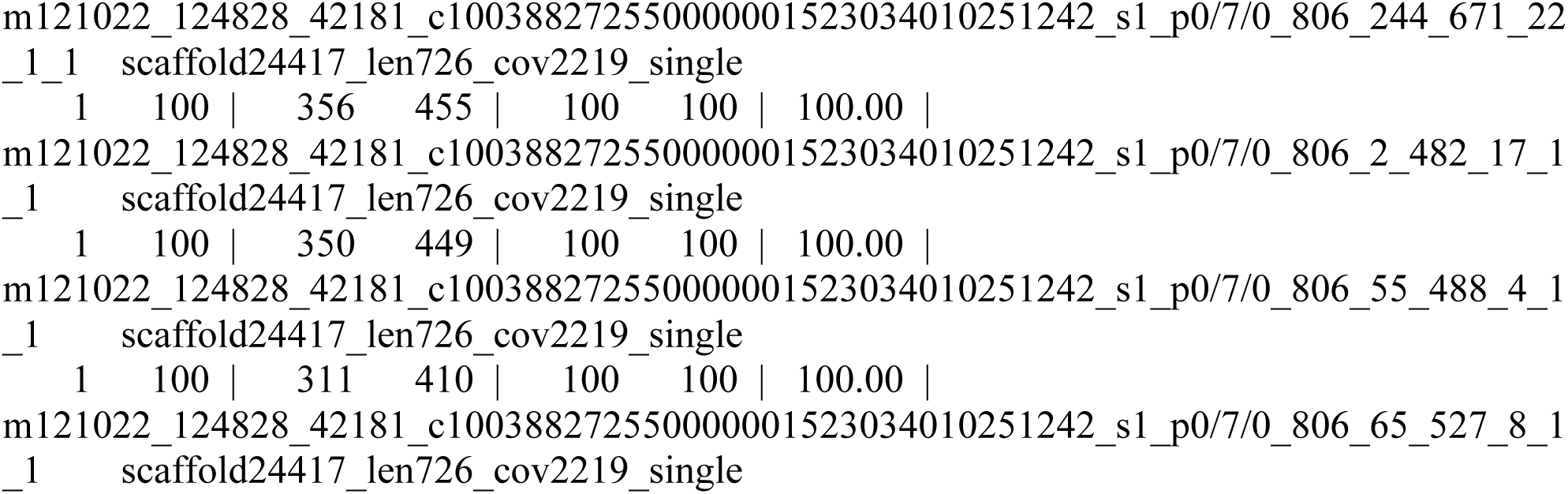
NUCMER based mapping of smaller Illumina reads coming from a single long PacBio read, to the assembly.

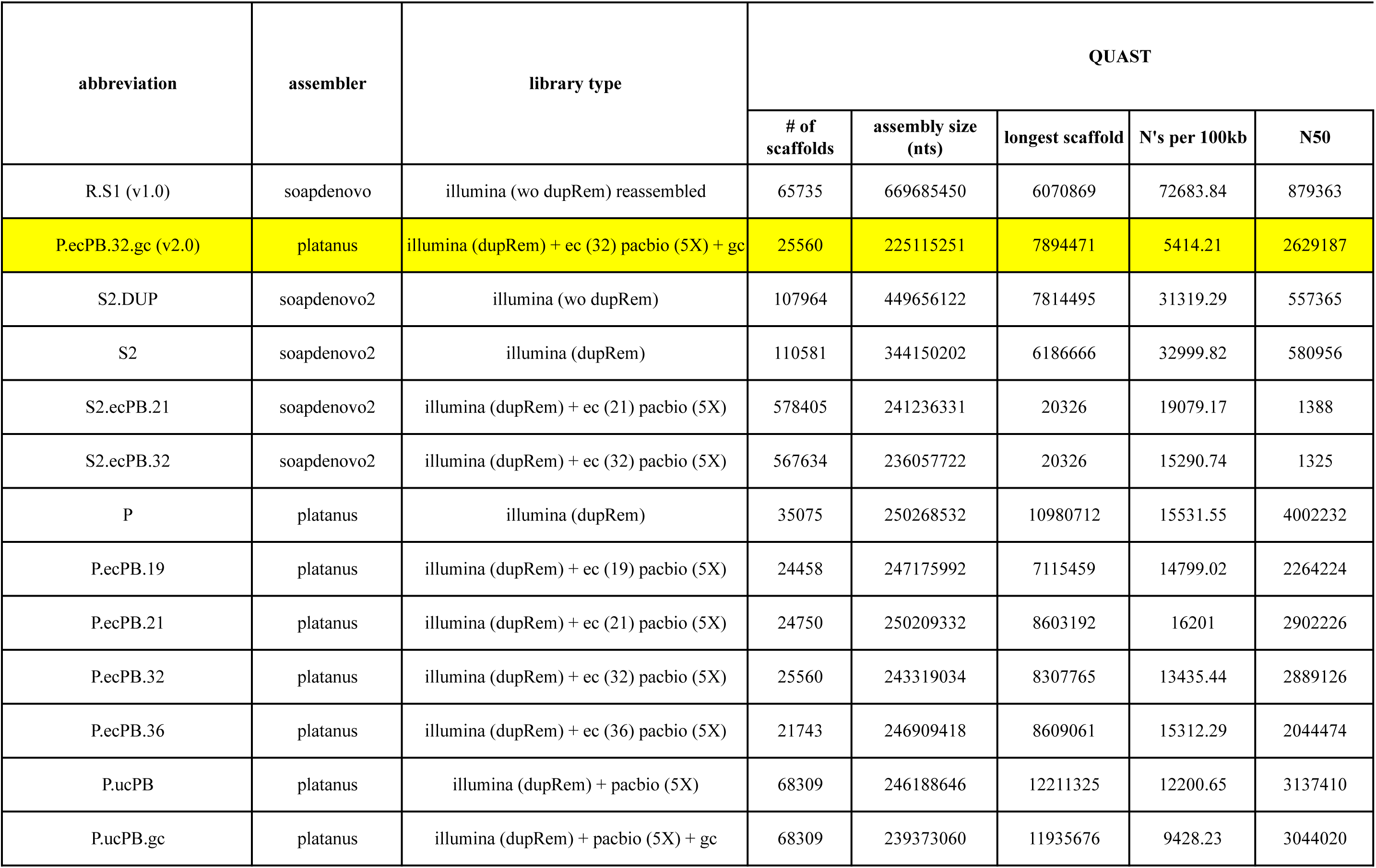

| abbreviation | assembler | library type | QUAST |  |  |  |  |
| --- | --- | --- | --- | --- | --- | --- | --- |
|  |  |  | # of scaffolds | assembly size (nts) | longest scaffold | N's per 100kb | N50 |
| R.S1 (v1.0) | soapdenovo | illumina (wo dupRem) reassembled | 65735 | 669685450 | 6070869 | 72683.84 | 879363 |
| P.ecPB.32.gc (v2.0) | platanus | illumina (dupRem) + ec (32) pacbio (5X) + gc | 25560 | 225115251 | 7894471 | 5414.21 | 2629187 |
| S2.DUP | soapdenovo2 | illumina (wo dupRem) | 107964 | 449656122 | 7814495 | 31319.29 | 557365 |
| S2 | soapdenovo2 | illumina (dupRem) | 110581 | 344150202 | 6186666 | 32999.82 | 580956 |
| S2.ecPB.21 | soapdenovo2 | illumina (dupRem) + ec (21) pacbio (5X) | 578405 | 241236331 | 20326 | 19079.17 | 1388 |
| S2.ecPB.32 | soapdenovo2 | illumina (dupRem) + ec (32) pacbio (5X) | 567634 | 236057722 | 20326 | 15290.74 | 1325 |
| P | platanus | illumina (dupRem) | 35075 | 250268532 | 10980712 | 15531.55 | 4002232 |
| P.ecPB.19 | platanus | illumina (dupRem) + ec (19) pacbio (5X) | 24458 | 247175992 | 7115459 | 14799.02 | 2264224 |
| P.ecPB.21 | platanus | illumina (dupRem) + ec (21) pacbio (5X) | 24750 | 250209332 | 8603192 | 16201 | 2902226 |
| P.ecPB.32 | platanus | illumina (dupRem) + ec (32) pacbio (5X) | 25560 | 243319034 | 8307765 | 13435.44 | 2889126 |
| P.ecPB.36 | platanus | illumina (dupRem) + ec (36) pacbio (5X) | 21743 | 246909418 | 8609061 | 15312.29 | 2044474 |
| P.ucPB | platanus | illumina (dupRem) + pacbio (5X) | 68309 | 246188646 | 12211325 | 12200.65 | 3137410 |
| P.ucPB.gc | platanus | illumina (dupRem) + pacbio (5X) + gc | 68309 | 239373060 | 11935676 | 9428.23 | 3044020 |

|  |  | NG50 (364 mb) | % covered<br>at 5X | % covered<br>at 10X | PASA (#s) |  |  | PASA (Lengths) |  |  | PASA (Exon<br>Lengths) |
| --- | --- | --- | --- | --- | --- | --- | --- | --- | --- | --- | --- |
| N75 | assembly size<br>(no gaps) |  |  |  | Valid<br>alignments | Failed<br>alignments | Assemblie<br>s | Valid<br>alignments | Failed alignments | Assemblies | Valid<br>alignments |
| 329406 | 182932349 | 1663167 | 13.66 | 6.83 | 61991 | 23523 | 52924 | 75719343 | 190033044 | 74739553 | 24479791 |
| 1149129 | 213036786 | 734691 | 65.50 | 32.75 | 77635 | 6584 | 61292 | 99453593 | 32030331 | 98694338 | 47829727 |
| 181445 | 308827017.1 | 836066 | 37.41 | 18.71 | 40416 | 43883 | 35880 | 21118152 | 214585399 | 20954880 | 9219905 |
| 194116 | 230581254.8 | 681660 | 32.88 | 16.44 | 72582 | 10851 | 58508 | 53795060 | 120907710 | 53154195 | 22605627 |
| 837 | 195210441.3 | 290 | 60.95 | 30.47 | 40082 | 26780 | 34723 | 11708648 | 13883902 | 11566814 | 9821552 |
| 821 | 199962749.5 | 270 | 62.38 | 31.19 | 40226 | 25494 | 34769 | 11794632 | 13941208 | 11647110 | 9899586 |
| 1489583 | 211397949.8 | 1587838 | 45.22 | 22.61 | 73800 | 11324 | 59057 | 85780130 | 53260936 | 85251475 | 41362113 |
| 876306 | 210596367.5 | 958049 | 59.64 | 29.82 | 70309 | 14030 | 56274 | 80711062 | 60735939 | 80164716 | 38639958 |
| 986022 | 209672918.1 | 1175333 | 58.76 | 29.38 | 69116 | 15189 | 55456 | 77277179 | 68307801 | 76849819 | 37102672 |
| 1243543 | 210628051.2 | 1112734 | 60.52 | 30.26 | 74576 | 8833 | 59412 | 91112744 | 40801767 | 90400020 | 43230588 |
| 973439 | 209101931.9 | 1030861 | 59.72 | 29.86 | 65304 | 19054 | 52778 | 66319838 | 80933035 | 65982106 | 31721203 |
| 1274166 | 216152031 | 1270061 | 60.16 | 30.08 | 73763 | 9728 | 58895 | 86665009 | 44526788 | 85994624 | 41389567 |
| 1252388 | 216804417.3 | 646875 | 61.87 | 30.94 | 73855 | 10533 | 58699 | 88744677 | 50211652 | 88084027 | 42629241 |

| abbreviation |  |  |  |  | predicted length = 1 nucleotide |  |  |  |  |  |  |
| --- | --- | --- | --- | --- | --- | --- | --- | --- | --- | --- | --- |
|  | GlimmerHMM (#s) |  | GlimmerHMM (Lengths) |  | GlimmerHMM (#s) |  | GlimmerHMM (Lengths) |  | PASA (#s) |  | PASA (I |
|  | Genes | Exons | Genes | Exons | Genes | Exons | Genes | Exons | Valid | Failed | Valid |
| R.S1 (v1.0) | 22816 | 59432 | 27884404 | 11837318 | 0 | 3 | 0 | 3 | 0 | 0 | 0 |
| P.ecPB.32.gc (v2.0) | 28553 | 128053 | 68723917 | 30727025 | 0 | 13 | 0 | 13 | 2 | 1 | 2 |
| S2.DUP | 34484 | 102919 | 54852164 | 20474483 | 0 | 12 | 0 | 12 | 1 | 9 | 1 |
| S2 | 32304 | 108656 | 60771387 | 23656610 | 0 | 5 | 0 | 5 | 1 | 2 | 1 |
| S2.ecPB.21 | 59707 | 90723 | 25691472 | 18635045 | 0 | 2 | 0 | 2 | 0 | 0 | 0 |
| S2.ecPB.32 | 60610 | 92091 | 26159705 | 19002432 | 0 | 1 | 0 | 1 | 0 | 0 | 0 |
| P | 25707 | 112438 | 60757663 | 25795802 | 0 | 9 | 0 | 9 | 2 | 2 | 2 |
| P.ecPB.19 | 26159 | 114069 | 62047674 | 26457349 | 0 | 5 | 0 | 5 | 2 | 1 | 2 |
| P.ecPB.21 | 25694 | 112341 | 60941486 | 25691804 | 0 | 13 | 0 | 13 | 2 | 6 | 2 |
| P.ecPB.32 | 25863 | 115978 | 62504333 | 26766180 | 0 | 13 | 0 | 13 | 4 | 1 | 4 |
| P.ecPB.36 | 24954 | 109392 | 58493707 | 23706845 | 0 | 7 | 0 | 7 | 1 | 2 | 1 |
| P.ucPB | 25911 | 115639 | 62033977 | 26174038 | 0 | 14 | 0 | 14 | 4 | 3 | 4 |
| P.ucPB.gc | 27356 | 121406 | 65264827 | 28187804 | 0 | 10 | 0 | 10 | 4 | 2 | 4 |

|  | predicted length <= 10 nucleotide |  |  |  |  |  |  |  |  |  |  |
| --- | --- | --- | --- | --- | --- | --- | --- | --- | --- | --- | --- |
| Lengths) | GlimmerHMM (#s) |  | GlimmerHMM (Lengths) |  | PASA (#s) |  | PASA (Lengths) |  | GlimmerHMM (#s) |  | GlimmerHM |
| Failed | Genes | Exons | Genes | Exons | Valid | Failed | Valid | Failed | Genes | Exons | Genes |
| 0 | 553 | 2012 | 3269 | 12424 | 451 | 724 | 2872 | 4524 | 1308 | 9296 | 22679 |
| 1 | 98 | 1599 | 673 | 9985 | 534 | 151 | 3351 | 965 | 286 | 11559 | 6071 |
| 9 | 526 | 2280 | 3066 | 14448 | 180 | 885 | 1188 | 5771 | 1349 | 13925 | 23681 |
| 2 | 522 | 2145 | 3051 | 13350 | 345 | 526 | 2169 | 3390 | 1311 | 12662 | 23602 |
| 0 | 1745 | 2232 | 10644 | 13682 | 90 | 128 | 588 | 801 | 7585 | 12001 | 181203 |
| 0 | 1721 | 2214 | 10485 | 13566 | 97 | 115 | 632 | 732 | 7635 | 12062 | 183275 |
| 2 | 211 | 1682 | 1251 | 10462 | 497 | 234 | 3176 | 1543 | 501 | 10644 | 8613 |
| 1 | 132 | 1612 | 760 | 9953 | 430 | 333 | 2711 | 2106 | 334 | 10414 | 6347 |
| 6 | 116 | 1584 | 682 | 9828 | 450 | 344 | 2782 | 2152 | 313 | 10314 | 5886 |
| 1 | 136 | 1572 | 812 | 9792 | 480 | 186 | 2993 | 1235 | 343 | 10715 | 6130 |
| 2 | 156 | 1667 | 890 | 10488 | 430 | 425 | 2708 | 2787 | 351 | 10949 | 5802 |
| 3 | 127 | 1646 | 778 | 10306 | 493 | 257 | 3070 | 1656 | 317 | 10986 | 5550 |
| 2 | 67 | 1602 | 381 | 9969 | 486 | 247 | 3026 | 1538 | 245 | 11165 | 5284 |

| predicted length <= 50 nucleotide |  |  |  |  | predicted length <= 100 nucleotide |  |  |  |  |  |  |  |
| --- | --- | --- | --- | --- | --- | --- | --- | --- | --- | --- | --- | --- |
| IM (Lengths) | PASA (#s) |  | PASA (Lengths) |  | GlimmerHMM (#s) |  | GlimmerHMM (Lengths) |  | PASA (#s) |  | PASA (Lengths) |  |
| Exons | Valid | Failed | Valid | Failed | Genes | Exons | Genes | Exons | Valid | Failed | Valid | Failed |
| 233878 | 3019 | 3875 | 84914 | 102181 | 1875 | 23993 | 65004 | 1352381 | 9540 | 11079 | 582283 | 651332 |
| 346117 | 5825 | 1210 | 195579 | 36596 | 478 | 45725 | 20609 | 2959090 | 31051 | 5248 | 2133009 | 345098 |
| 384755 | 1529 | 7424 | 46934 | 223183 | 1925 | 40351 | 66008 | 2386791 | 5753 | 26156 | 369403 | 1646989 |
| 351635 | 3358 | 4682 | 106802 | 142899 | 1957 | 40903 | 71841 | 2497568 | 14455 | 18716 | 954945 | 1214213 |
| 319161 | 962 | 1143 | 30716 | 36185 | 15636 | 34432 | 793299 | 2036255 | 4417 | 5570 | 297024 | 379502 |
| 321267 | 950 | 1156 | 30010 | 37114 | 15752 | 34772 | 800883 | 2060085 | 4389 | 5650 | 295320 | 385927 |
| 311165 | 5117 | 1906 | 169515 | 57635 | 676 | 40990 | 21659 | 2632658 | 26056 | 8056 | 1776258 | 528900 |
| 306500 | 4761 | 2398 | 159521 | 72508 | 486 | 40778 | 18170 | 2631236 | 24833 | 10141 | 1701956 | 664971 |
| 303668 | 4646 | 2621 | 153767 | 79546 | 449 | 40713 | 16357 | 2630525 | 24304 | 10920 | 1663611 | 714191 |
| 318217 | 5286 | 1603 | 177301 | 48875 | 503 | 42146 | 18052 | 2723607 | 27911 | 6710 | 1914064 | 440339 |
| 321218 | 4368 | 3379 | 145263 | 102697 | 486 | 41038 | 15720 | 2620577 | 21680 | 14105 | 1473488 | 924155 |
| 323189 | 5267 | 1913 | 175573 | 56968 | 453 | 42407 | 15753 | 2727163 | 26973 | 7970 | 1842651 | 520660 |
| 332259 | 5305 | 1937 | 178157 | 58931 | 407 | 43702 | 17516 | 2822003 | 27776 | 8299 | 1904545 | 545248 |

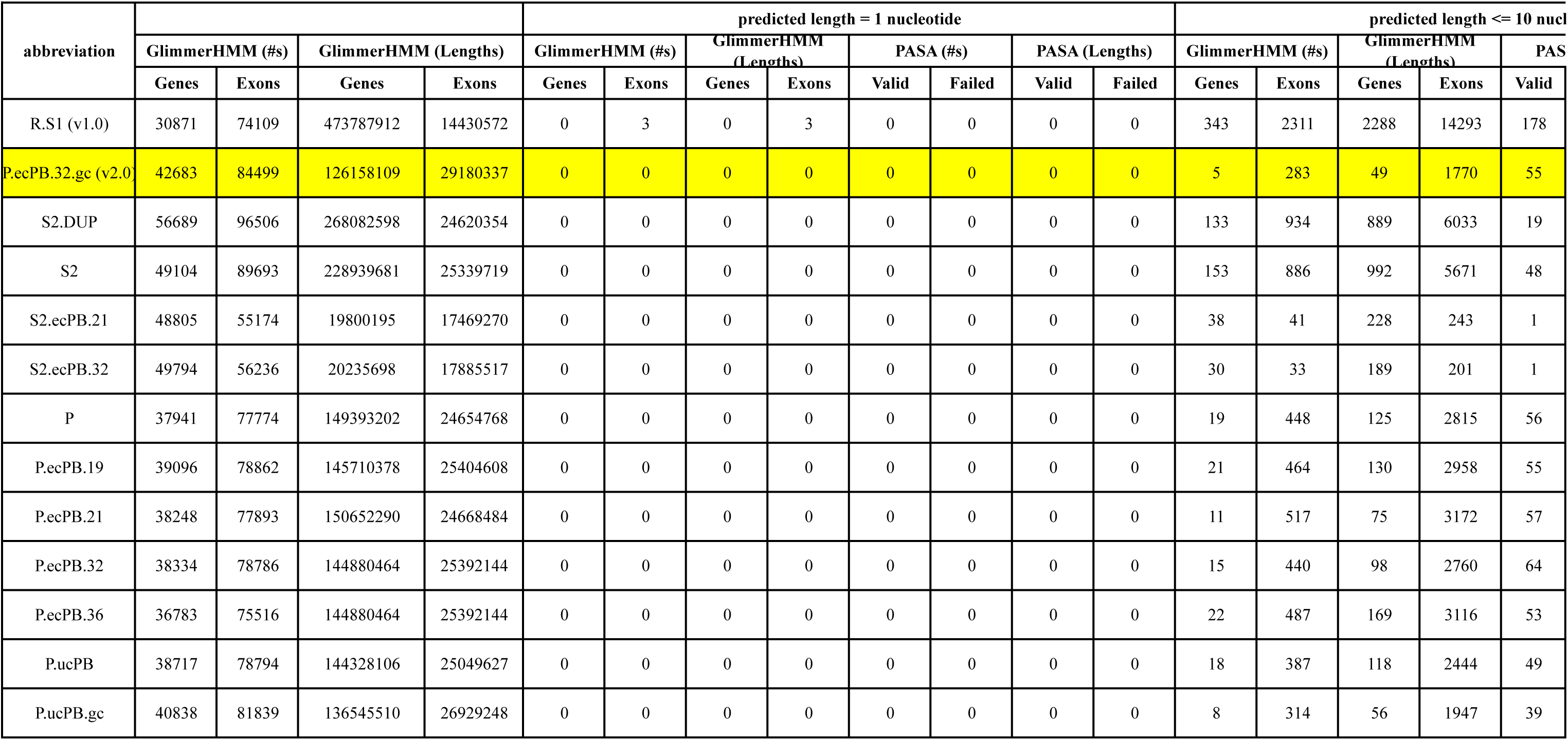

| abbreviation |  |  |  |  | predicted length = 1 nucleotide |  |  |  |  |  |  |  | predicted length <= 10 nucl |  |  |  |  |
| --- | --- | --- | --- | --- | --- | --- | --- | --- | --- | --- | --- | --- | --- | --- | --- | --- | --- |
|  | GlimmerHMM (#s) |  | GlimmerHMM (Lengths) |  | GlimmerHMM (#s) |  | GlimmerHMM (Lengths) |  | PASA (#s) |  | PASA (Lengths) |  | GlimmerHMM (#s) |  | GlimmerHMM (Lengths) |  | PAS |
|  | Genes | Exons | Genes | Exons | Genes | Exons | Genes | Exons | Valid | Failed | Valid | Failed | Genes | Exons | Genes | Exons | Valid |
| R.S1 (v1.0) | 30871 | 74109 | 473787912 | 14430572 | 0 | 3 | 0 | 3 | 0 | 0 | 0 | 0 | 343 | 2311 | 2288 | 14293 | 178 |
| P.ecPB.32.gc (v2.0) | 42683 | 84499 | 126158109 | 29180337 | 0 | 0 | 0 | 0 | 0 | 0 | 0 | 0 | 5 | 283 | 49 | 1770 | 55 |
| S2.DUP | 56689 | 96506 | 268082598 | 24620354 | 0 | 0 | 0 | 0 | 0 | 0 | 0 | 0 | 133 | 934 | 889 | 6033 | 19 |
| S2 | 49104 | 89693 | 228939681 | 25339719 | 0 | 0 | 0 | 0 | 0 | 0 | 0 | 0 | 153 | 886 | 992 | 5671 | 48 |
| S2.ecPB.21 | 48805 | 55174 | 19800195 | 17469270 | 0 | 0 | 0 | 0 | 0 | 0 | 0 | 0 | 38 | 41 | 228 | 243 | 1 |
| S2.ecPB.32 | 49794 | 56236 | 20235698 | 17885517 | 0 | 0 | 0 | 0 | 0 | 0 | 0 | 0 | 30 | 33 | 189 | 201 | 1 |
| P | 37941 | 77774 | 149393202 | 24654768 | 0 | 0 | 0 | 0 | 0 | 0 | 0 | 0 | 19 | 448 | 125 | 2815 | 56 |
| P.ecPB.19 | 39096 | 78862 | 145710378 | 25404608 | 0 | 0 | 0 | 0 | 0 | 0 | 0 | 0 | 21 | 464 | 130 | 2958 | 55 |
| P.ecPB.21 | 38248 | 77893 | 150652290 | 24668484 | 0 | 0 | 0 | 0 | 0 | 0 | 0 | 0 | 11 | 517 | 75 | 3172 | 57 |
| P.ecPB.32 | 38334 | 78786 | 144880464 | 25392144 | 0 | 0 | 0 | 0 | 0 | 0 | 0 | 0 | 15 | 440 | 98 | 2760 | 64 |
| P.ecPB.36 | 36783 | 75516 | 144880464 | 25392144 | 0 | 0 | 0 | 0 | 0 | 0 | 0 | 0 | 22 | 487 | 169 | 3116 | 53 |
| P.ucPB | 38717 | 78794 | 144328106 | 25049627 | 0 | 0 | 0 | 0 | 0 | 0 | 0 | 0 | 18 | 387 | 118 | 2444 | 49 |
| P.ucPB.gc | 40838 | 81839 | 136545510 | 26929248 | 0 | 0 | 0 | 0 | 0 | 0 | 0 | 0 | 8 | 314 | 56 | 1947 | 39 |

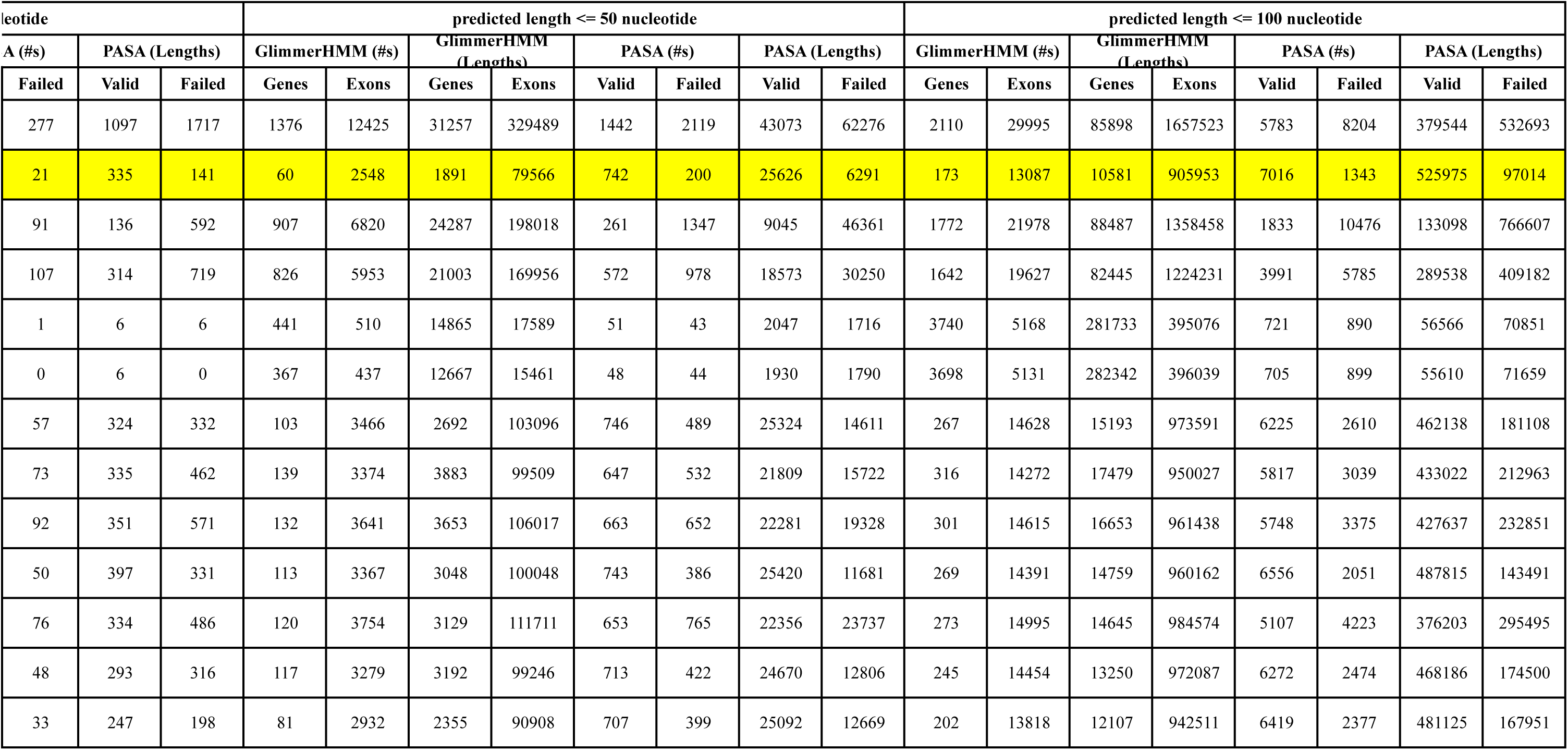

| abbreviation |  |  |  |  | predicted length = 1 nucleotide |  |  |  |  |  |  |  | predicted length <= 10 nucleotide |  |  |  |  |  |  |
| --- | --- | --- | --- | --- | --- | --- | --- | --- | --- | --- | --- | --- | --- | --- | --- | --- | --- | --- | --- |
|  | GlimmerHMM (#s) |  | GlimmerHMM (Lengths) |  | GlimmerHMM (#s) |  | GlimmerHMM (Lengths) |  | PASA (#s) |  | PASA (Lengths) |  | GlimmerHMM (#s) |  | GlimmerHMM (Lengths) |  | PASA (#s) |  | PASA (L |
|  | Genes | Exons | Genes | Exons | Genes | Exons | Genes | Exons | Valid | Failed | Valid | Failed | Genes | Exons | Genes | Exons | Valid | Failed | Valid |
| R.S1 (v1.0) | 54500 | 97841 | 4.3E+08 | 1.8E+07 | 148 | 185 | 148 | 185 | 0 | 0 | 0 | 0 | 603 | 3464 | 3177 | 20146 | 243 | 459 | 1555 |
| P.ecPB.32.gc (v2.0) | 78021 | 115562 | 1.1E+08 | 3.6E+07 | 0 | 66 | 0 | 66 | 1 | 0 | 1 | 0 | 66 | 556 | 130 | 3057 | 120 | 47 | 762 |
| S2.DUP | 114527 | 153380 | 2.3E+08 | 3.7E+07 | 139 | 179 | 139 | 179 | 0 | 0 | 0 | 0 | 303 | 1623 | 1244 | 9141 | 27 | 200 | 174 |
| S2 | 93737 | 132158 | 2E+08 | 3.5E+07 | 169 | 204 | 169 | 204 | 0 | 0 | 0 | 0 | 340 | 1608 | 1329 | 8958 | 117 | 232 | 751 |
| S2.ecPB.21 | 77294 | 84692 | 2.5E+07 | 2.3E+07 | 559 | 568 | 559 | 568 | 1 | 3 | 1 | 3 | 661 | 689 | 1274 | 1403 | 10 | 9 | 61 |
| S2.ecPB.32 | 78991 | 86476 | 2.6E+07 | 2.3E+07 | 544 | 555 | 544 | 555 | 1 | 2 | 1 | 2 | 627 | 657 | 1119 | 1250 | 11 | 10 | 71 |
| P | 69929 | 106381 | 1.3E+08 | 3.1E+07 | 134 | 140 | 134 | 140 | 0 | 0 | 0 | 0 | 163 | 956 | 328 | 5125 | 109 | 124 | 682 |
| P.ecPB.19 | 71427 | 107756 | 1.2E+08 | 3.1E+07 | 45 | 51 | 45 | 51 | 0 | 0 | 0 | 0 | 67 | 869 | 207 | 4990 | 118 | 130 | 728 |
| P.ecPB.21 | 70084 | 106364 | 1.3E+08 | 3.1E+07 | 50 | 64 | 50 | 64 | 0 | 0 | 0 | 0 | 79 | 1000 | 257 | 5841 | 121 | 150 | 764 |
| P.ecPB.32 | 70608 | 107431 | 1.2E+08 | 3.1E+07 | 76 | 85 | 76 | 85 | 1 | 0 | 1 | 0 | 100 | 868 | 234 | 5001 | 134 | 82 | 864 |
| P.ecPB.36 | 67855 | 103858 | 1.3E+08 | 2.9E+07 | 45 | 52 | 45 | 52 | 0 | 1 | 0 | 1 | 71 | 937 | 231 | 5596 | 100 | 172 | 613 |
| P.ucPB | 72000 | 108702 | 1.2E+08 | 3.1E+07 | 67 | 76 | 67 | 76 | 0 | 1 | 0 | 1 | 92 | 833 | 247 | 4734 | 129 | 90 | 798 |
| P.ucPB.gc | 75114 | 112381 | 1.2E+08 | 3.3E+07 | 33 | 44 | 33 | 44 | 0 | 1 | 0 | 1 | 43 | 677 | 107 | 3948 | 119 | 77 | 736 |

|  | predicted length <= 50 nucleotide |  |  |  |  |  |  |  | predicted length <= 100 nucleotide |  |  |  |  |  |  |  |
| --- | --- | --- | --- | --- | --- | --- | --- | --- | --- | --- | --- | --- | --- | --- | --- | --- |
| Lengths) | GlimmerHMM (#s) |  | GlimmerHMM (Lengths) |  | PASA (#s) |  | PASA (Lengths) |  | GlimmerHMM (#s) |  | GlimmerHMM (Lengths) |  | PASA (#s) |  | PASA (Lengths) |  |
| Failed | Genes | Exons | Genes | Exons | Valid | Failed | Valid | Failed | Genes | Exons | Genes | Exons | Valid | Failed | Valid | Failed |
| 2862 | 1862 | 16063 | 37773 | 407084 | 1901 | 2914 | 57170 | 83268 | 2788 | 37081 | 107490 | 2005806 | 7108 | 9993 | 460903 | 630127 |
| 302 | 141 | 3857 | 2544 | 115861 | 1237 | 325 | 41353 | 9807 | 340 | 17382 | 18369 | 1170784 | 9021 | 1705 | 658050 | 118323 |
| 1238 | 1163 | 9224 | 27025 | 255843 | 366 | 2124 | 12203 | 70975 | 2338 | 28647 | 115491 | 1749508 | 2286 | 12847 | 164734 | 915748 |
| 1467 | 1212 | 8279 | 27355 | 225200 | 946 | 1589 | 29541 | 48212 | 2319 | 25696 | 111480 | 1567929 | 5220 | 7549 | 366574 | 517553 |
| 40 | 1985 | 2207 | 48461 | 56113 | 142 | 152 | 5194 | 5416 | 9307 | 11896 | 629479 | 829282 | 1306 | 1604 | 99113 | 123559 |
| 48 | 1917 | 2148 | 47265 | 55149 | 137 | 159 | 4880 | 5706 | 9263 | 11877 | 630691 | 831311 | 1301 | 1618 | 98816 | 124031 |
| 770 | 317 | 5252 | 4694 | 146697 | 1205 | 848 | 40134 | 24675 | 567 | 19164 | 23802 | 1227977 | 7908 | 3455 | 571511 | 228602 |
| 797 | 213 | 5058 | 4752 | 143087 | 1132 | 861 | 37029 | 24857 | 457 | 18710 | 23691 | 1206458 | 7470 | 3916 | 540496 | 265242 |
| 977 | 193 | 5252 | 3544 | 146627 | 1043 | 975 | 34482 | 28504 | 429 | 19104 | 21680 | 1223274 | 7262 | 4338 | 527553 | 291587 |
| 510 | 232 | 4992 | 4315 | 142129 | 1222 | 577 | 39916 | 17371 | 462 | 18903 | 22056 | 1223364 | 8409 | 2604 | 608198 | 176624 |
| 1048 | 213 | 5463 | 4430 | 154569 | 1005 | 1256 | 33439 | 37757 | 435 | 19666 | 21462 | 1259239 | 6552 | 5532 | 473329 | 372864 |
| 575 | 233 | 4982 | 4334 | 141844 | 1191 | 647 | 39052 | 19491 | 470 | 19017 | 23024 | 1233823 | 8027 | 3135 | 581825 | 213970 |
| 461 | 132 | 4421 | 2984 | 129634 | 1156 | 604 | 38194 | 17946 | 370 | 18274 | 22125 | 1209139 | 8195 | 3081 | 596091 | 212526 |

**Supplementary Table 3.** Repeat element classification

| Repeat category | Sub-category | Number | Content (bases) | Content (% of genome assembly size) |
| --- | --- | --- | --- | --- |
| ***Retrotransposons*** | *LTR retrotransposon* |  |  |  |
|  | Gypsy | 16983 | 9699390 | 4.31 |
|  | Copia | 14183 | 9031714 | 4.01 |
|  | Caulimovirus | 1298 | 919770 | 0.41 |
|  | Unclassified | 19 | 8004 | 0.004 |
|  | *Non-LTR retrotransposon* |  |  |  |
|  | LINEs | 2428 | 845972 | 0.38 |
|  | SINEs | 0 | 0 | 0 |
| ***DNA Transposons*** | *Terminal Inverted Repeats (TIRs)* |  |  |  |
|  | MuLE-MuDR | 6261 | 2797241 | 1.24 |
|  | hAT | 1776 | 833096 | 0.37 |
|  | PIF/Harbinger | 407 | 152389 | 0.07 |
|  | En-Spm | 2143 | 969277 | 0.43 |
|  | others | 80 | 4328 | 0.002 |
| ***MITEs*** |  | 3782 | 707942 | 0.31 |
| ***RC/Helitron*** |  | 173 | 11906 | 0.005 |
| ***Unclassified Sequences*** |  | 98680 | 30568122 | 13.58 |
| ***Simple repeats*** |  | 1297 | 284765 | 0.13 |
| **Non redundant Repeat Content** |  | N.A. | **54375206** | **24.15** |

**Supplementary File 2.**
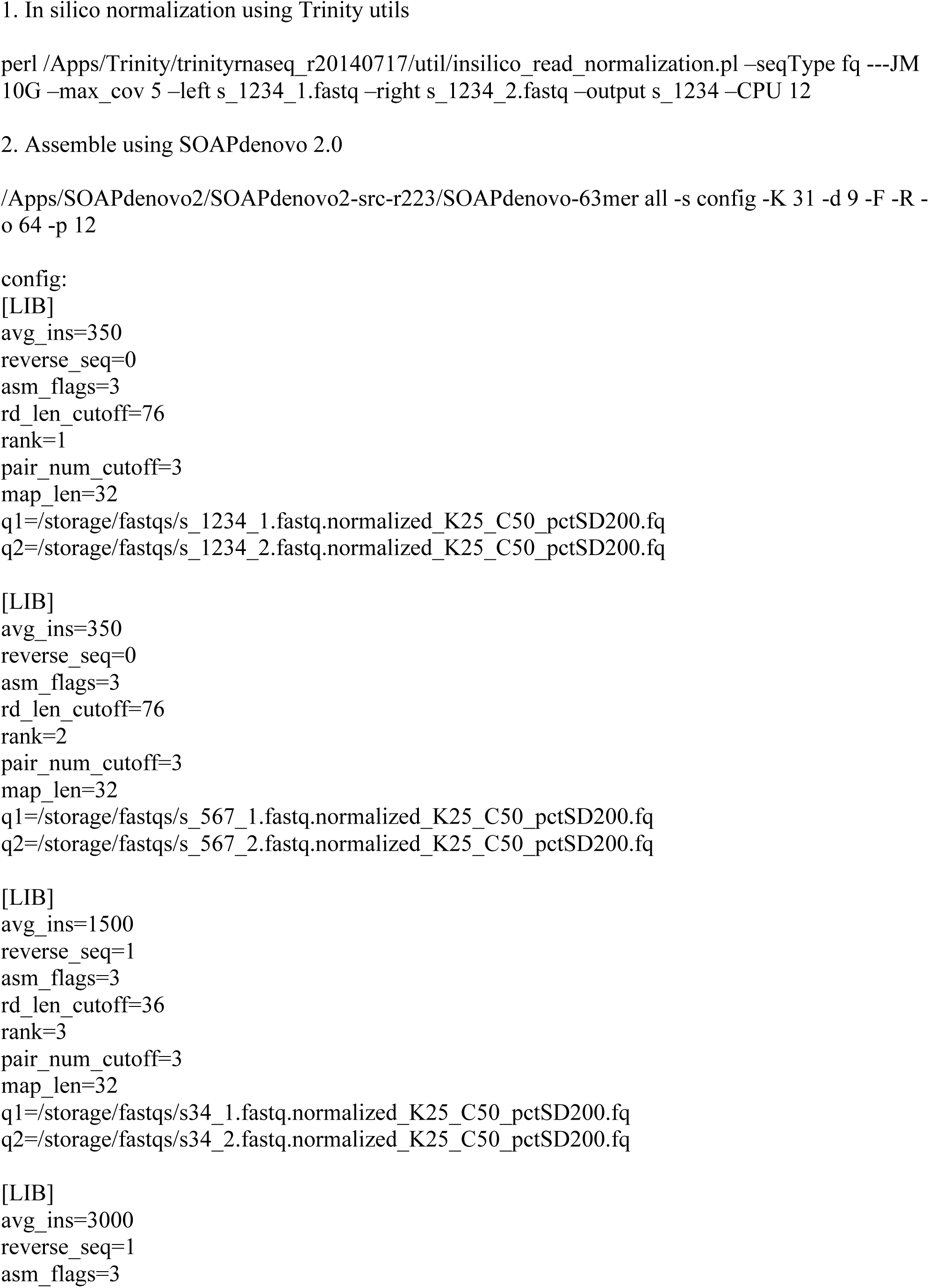
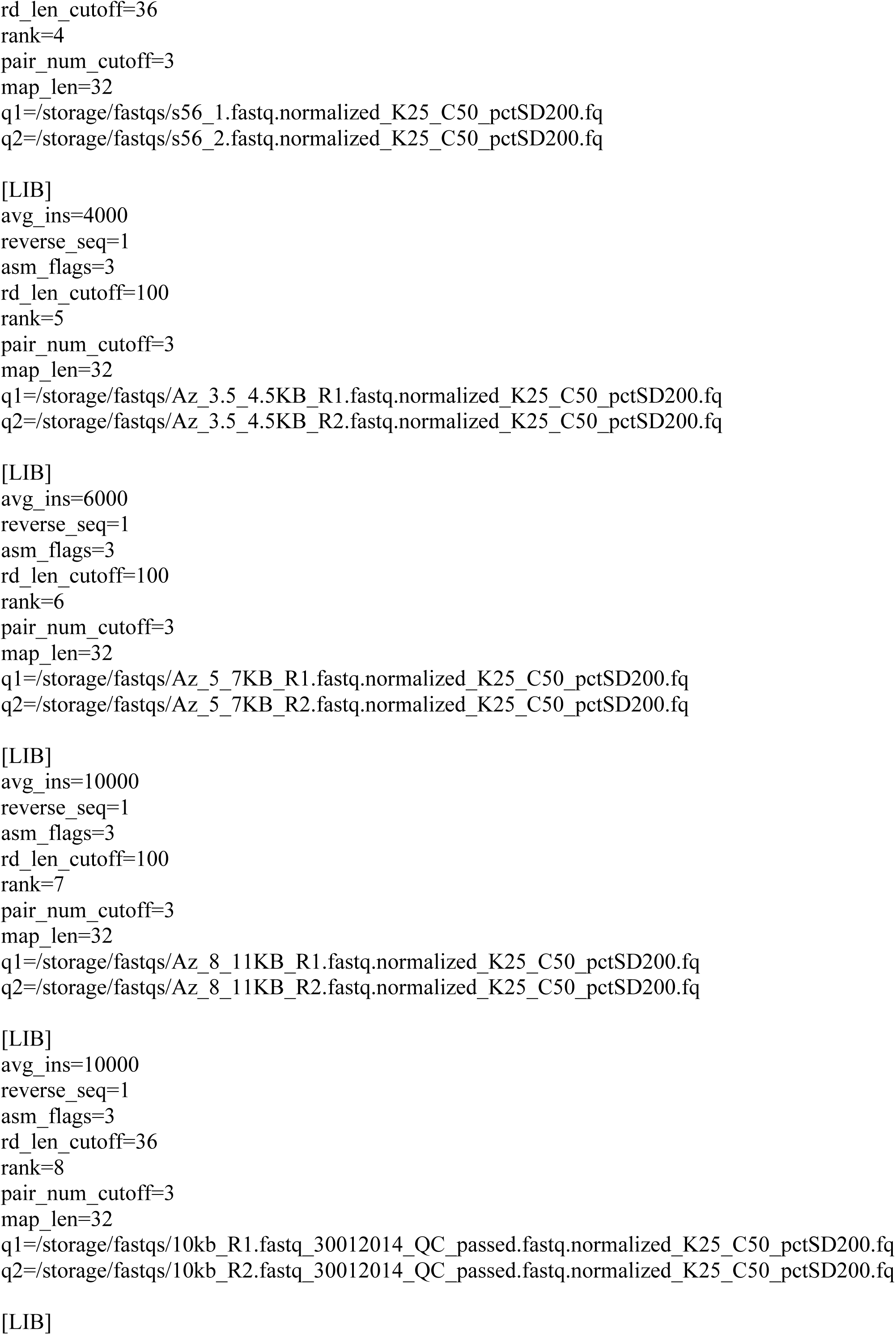
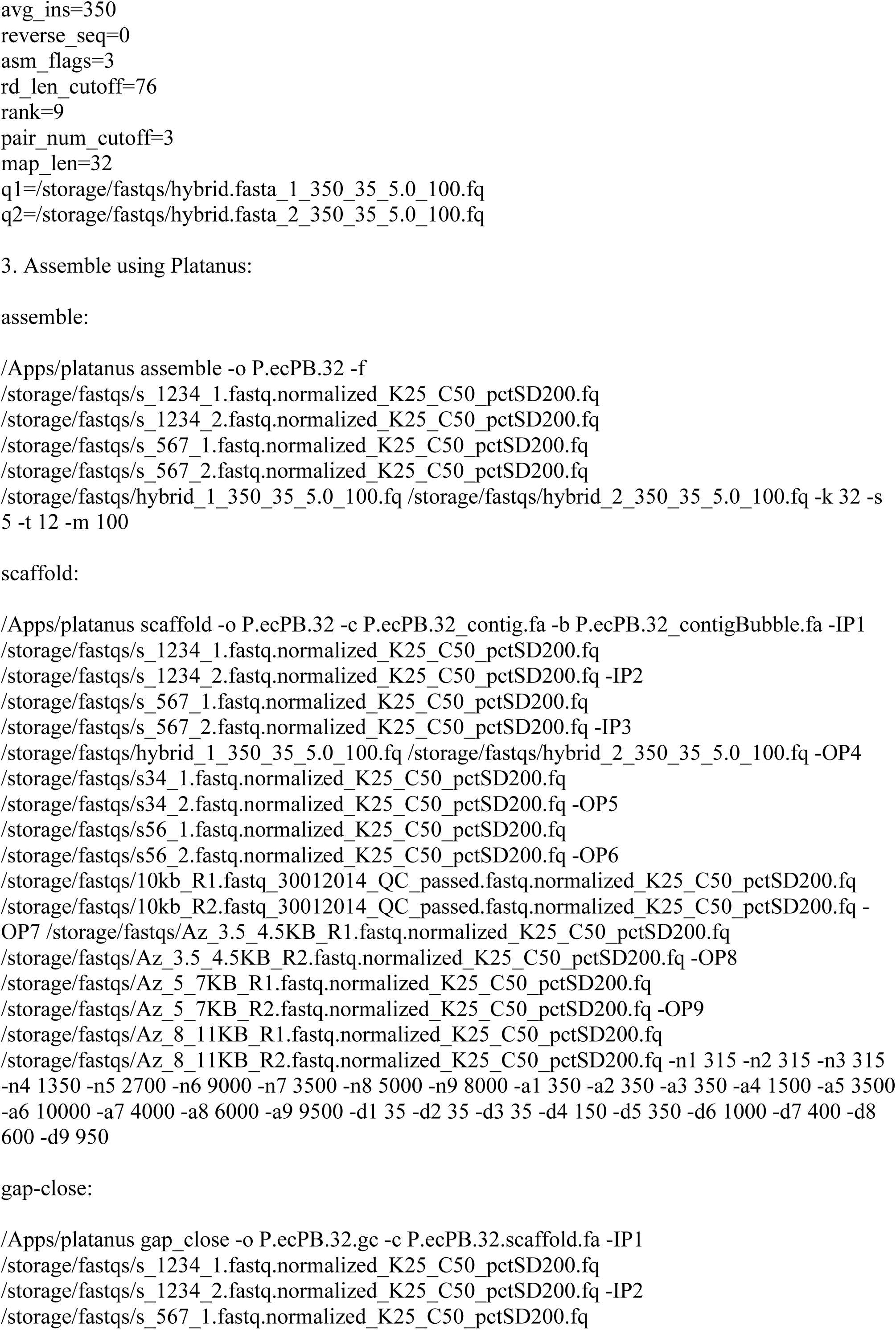
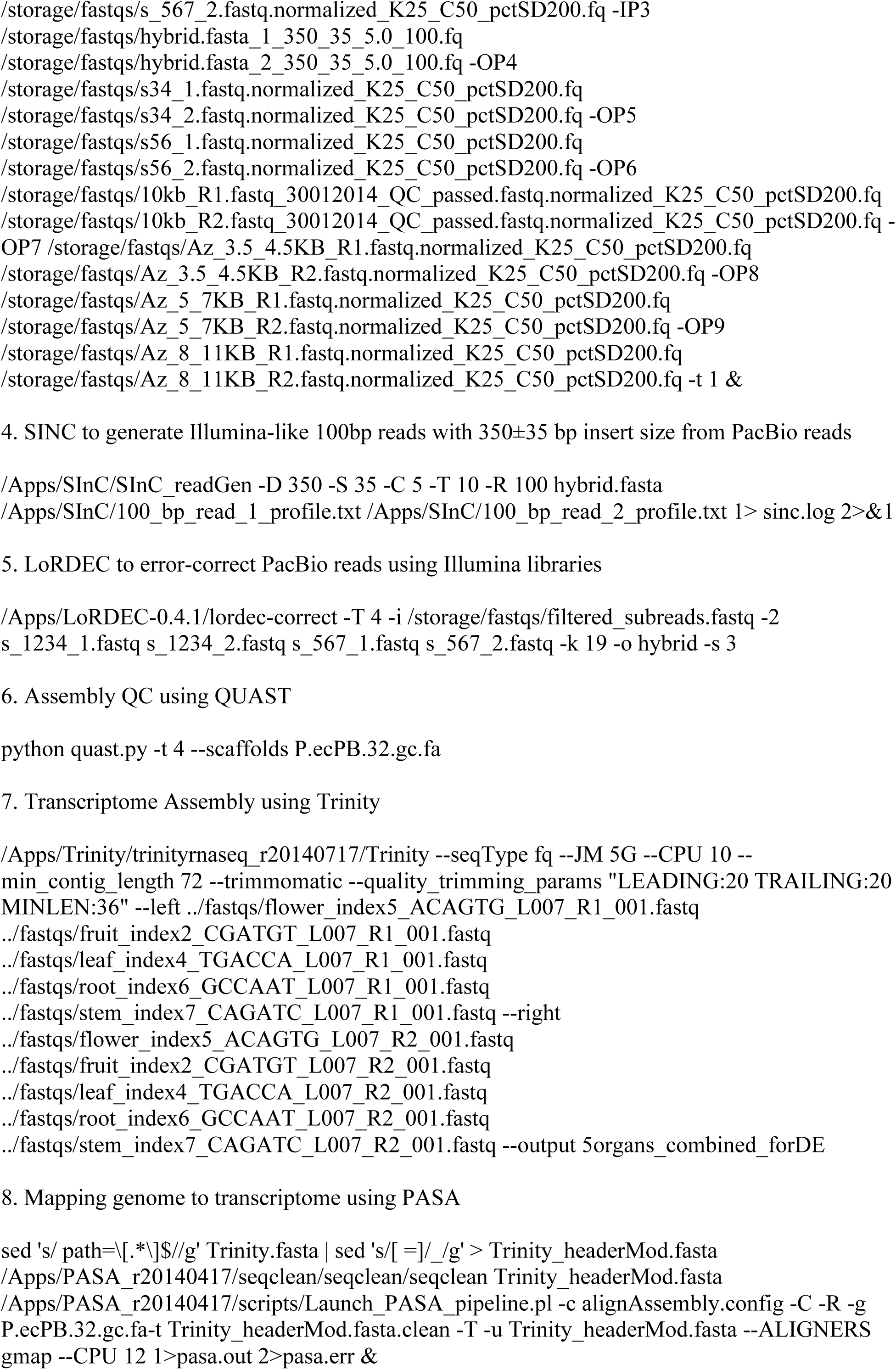
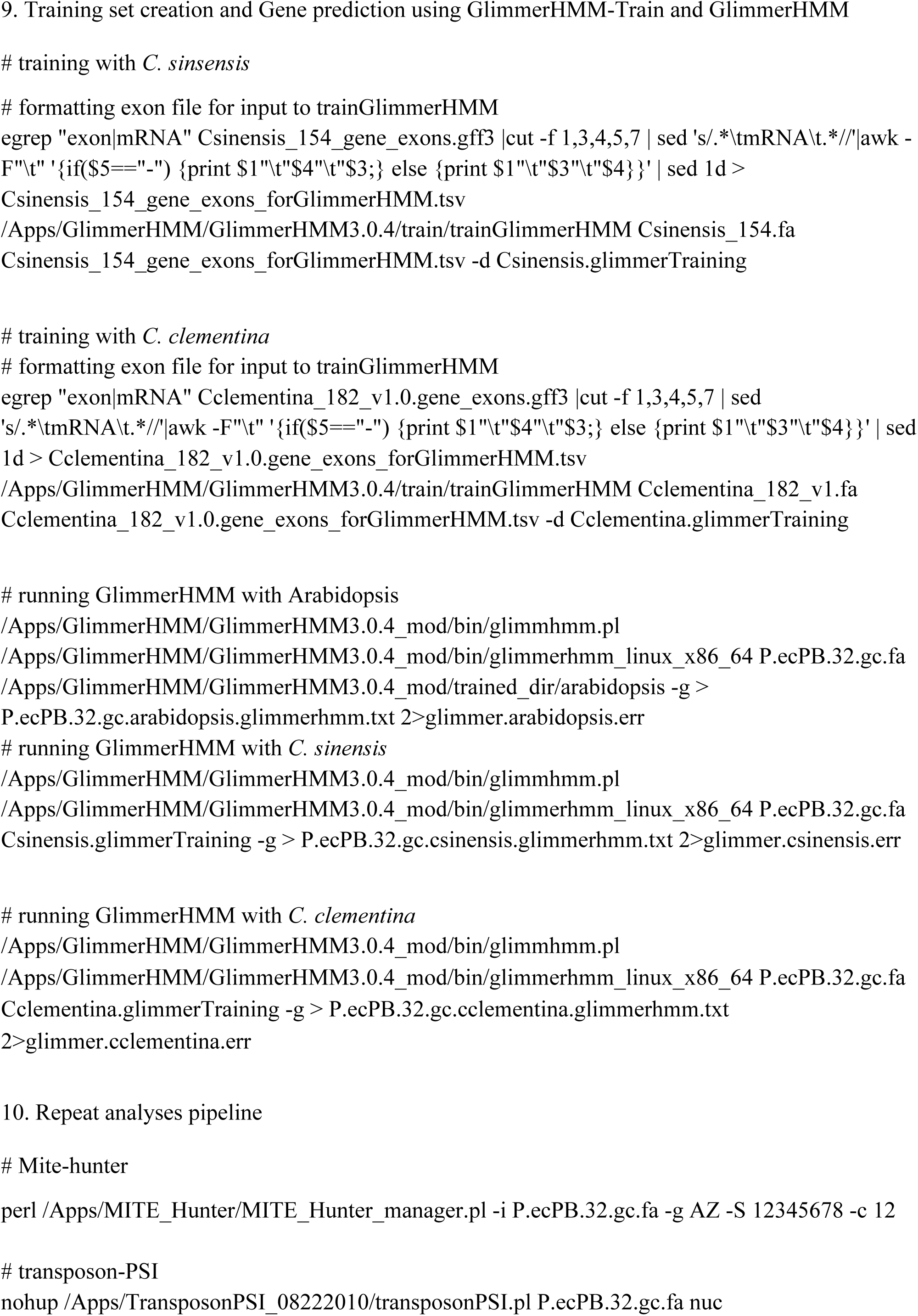
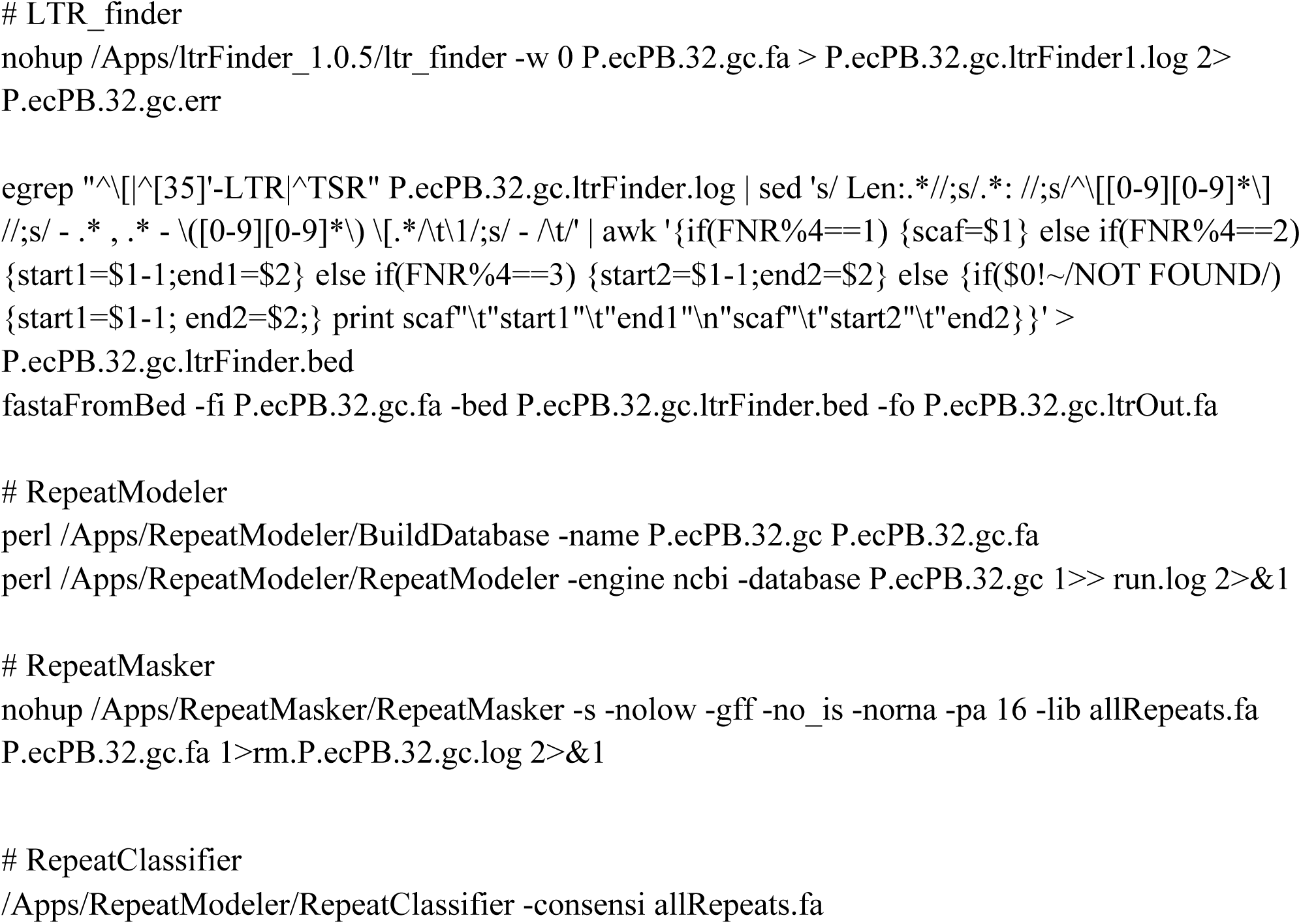
Scripts used in the study

